# Enhancing CO_2_-valorization using *Clostridium autoethanogenum* for sustainable fuel and chemicals production

**DOI:** 10.1101/2020.01.23.917666

**Authors:** James K. Heffernan, Kaspar Valgepea, Renato de Souza Pinto Lemgruber, Isabella Casini, Manuel Plan, Ryan Tappel, Sean D. Simpson, Michael Köpke, Lars K. Nielsen, Esteban Marcellin

**Affiliations:** Australian Institute for Bioengineering and Nanotechnology, The University of Queensland, Building 75, Corner College Rd & Cooper Rd, St. Lucia 4067, QLD, Australia; ERA Chair in Gas Fermentation Technologies, Institute of Technology, University of Tartu, Nooruse 1, 50411 Tartu, Estonia; Center for Applied Geosciences, University of Tübingen, Hölderlinstr. 12, 72074, Tübingen, Germany; Queensland Node of Metabolomics Australia, The University of Queensland, Building 75, Corner College Rd & Cooper Rd, St. Lucia 4067, QLD, Australia; LanzaTech Inc., 8045 Lamon Ave Suite 400, Skokie, IL 60077, USA

**Keywords:** gas fermentation, *Clostridium autoethanogenum*, carbon dioxide, valorization, carbon recycling, fuel and chemical platforms

## Abstract

Acetogenic bacteria can convert waste gases into fuels and chemicals. Design of bioprocesses for waste carbon valorization requires quantification of steady-state carbon flows. Here, steady-state quantification of autotrophic chemostats containing *Clostridium autoethanogenum* grown on CO_2_ and H_2_ revealed that captured carbon (460 ± 80 mmol/gDCW/day) had a significant distribution to ethanol (54 ± 3 mol% with a 2.4 ± 0.3 g/L titer). We were impressed with this initial result, but also observed limitations to biomass concentration and growth rate. Metabolic modelling predicted culture performance and indicated significant metabolic adjustments when compared to fermentation with CO as the carbon source. Moreover, modelling highlighted flux to pyruvate, and subsequently reduced ferredoxin, as a target for improving CO_2_ and H_2_ fermentation. Supplementation with a small amount of CO enabled co-utilisation with CO_2_, and enhanced CO_2_ fermentation performance significantly, while maintaining an industrially relevant product profile. Additionally, the highest specific flux through the Wood-Ljungdahl pathway was observed during co-utilization of CO_2_ and CO. Furthermore, the addition of CO led to superior CO_2_-valorizing characteristics (9.7 ± 0.4 g/L ethanol with a 66 ± 2 mol% distribution, and 540 ± 20 mmol CO_2_/gDCW/day). Similar industrial processes are commercial or currently being scaled up, indicating CO-supplemented CO_2_ and H_2_ fermentation has high potential for sustainable fuel and chemical production. This work also provides a reference dataset to advance our understanding of CO_2_ gas fermentation, which can contribute to mitigating climate change.

## Introduction

Gas fermentation has attractive waste carbon valorization properties, for which the need is intensifying (Emerson and Stephanopoulos, 2019; IPCC, 2014). Recently, LanzaTech commercialized the first waste gas-to-ethanol process, efficiently incorporating the carbon from steel mill off-gas into fuel quality ethanol *via* the model acetogen *Clostridium autoethanogenum*. The key carbon source — carbon monoxide (CO) — accounts for a significant portion of steel mill off-gas and synthesis gas (syngas), which can be generated from multiple high-volume, non-gaseous waste feedstocks (e.g. biomass, municipal solid waste) (Liew et al., 2016). Therefore, LanzaTech’s process is significant in that it valorizes waste carbon by fusing two one-carbon gas molecules (C1) into liquid fuel. Furthermore, Handler et al. (2016) found that ethanol produced by LanzaTech’s process reduced greenhouse gas emissions by 67 to 98% when compared to petroleum gasoline on an energy content and “cradle-to-grave” basis (feedstock dependent). Carbon dioxide (CO_2_) represents a more diverse and plentiful waste stream compared to CO (International Panel on Climate Change (IPCC), 2014), thus embodying a feedstock with greater climate change mitigation and carbon recycling potential.

Increasing acetogenic carbon capture as CO_2_ would build on the success of commercial gas fermentation and continue the expansion of the technology as a platform for sustainable chemical production (Bengelsdorf et al., 2018; Müller, 2019; Redl et al., 2017). Compared to other CO_2_ valorization methods, acetogens are ideal candidates due to their high metabolic efficiency, ability to handle variable gas compositions, high product specificity, scalability, and low susceptibility to poisoning by sulphur, chlorine, and tars (Artz et al., 2018; Liew et al., 2016). However, metabolism of CO_2_ requires an energy source, for which some see an appropriate solution is lacking (Emerson and Stephanopoulos, 2019).

Gas fermenting acetogens harbor the Wood-Ljungdahl pathway (WLP) (Drake et al., 2008), a non-photosynthetic C1-fixation metabolic pathway with the highest-known theoretical thermodynamic efficiency (Fast and Papoutsakis, 2012; Müller, 2019; Schuchmann and Müller, 2014). Various potential energy sources exist for metabolizing CO_2_, primarily hydrogen, nitrates, sugars, and arginine. Yet, acetogenic CO_2_ valorization, which is actively being developed for industrial implementation (Tizard and Sechrist, 2015), poses challenges along with promise. These include potential adenosine triphosphate (ATP) starvation in autotrophic conditions and carbon catabolite repression in hetero/mixotrophic conditions (Emerson and Stephanopoulos, 2019).

Hydrogen (H_2_) is the most recognized energy source for CO_2_ utilization — as metabolism of sugars or nitrates cause shifts in metabolism that result in lower CO_2_ or H_2_ utilization (Emerson and Stephanopoulos, 2019 & Liew et al., 2016). H_2_ production will also logically transition to renewable sources in the future, whereas production of sugars and nitrates are dependent on less-sustainable methods. Furthermore, levelized cost predictions for solar H_2_ indicate a 30% reduction by 2030, potentially becoming competitive with the current levelized cost of fossil fuel derived H_2_ by 2035 (Detz et al., 2018; Glenk and Reichelstein, 2019). This is in part due to rapidly decreasing solar electricity costs (IRENA, 2017) and projections of H_2_ electrolysis technology development (Detz et al., 2018; Glenk and Reichelstein, 2019). Similarly, atmospheric CO_2_ capture *via* direct air contact showed promising feasibility recently (Keith et al., 2018), which represents an essential development for carbon recycling (Otto et al., 2015). Various power-to-gas technologies are being discussed for mediating fluctuations in renewable power generation (Götz et al., 2016). By extension, gas fermentation to liquid products could couple mediation of renewable power fluctuations to carbon recycling (Redl et al., 2017). This provides an attractive new opportunity for bacterial artificial-photosynthesis, whereby renewable H_2_ supplementation facilitates acetogenic CO_2_ valorization (Claassens et al., 2016; Haas et al., 2018).

Continuous culture bioprocesses are preferable to batch or fed-batch fermentation bioprocesses (Hoskisson and Hobbs, 2005). Furthermore, systems-level quantification is essential for design-build-test-learn bioprocess optimization by metabolic engineering (Valgepea et al., 2017). Therefore, obtaining quantitative datasets from steady-state chemostat cultures, whose analyses are comparable between experiments, is important for development of these systems (Adamberg et al., 2015). Whilst Bengelsdorf et al. (2018) reviewed autotrophic acetogen growth on CO_2_ and H_2_ (CO_2_+H_2_), and Mock et al. (2015) provided notable insight into the CO_2_+H_2_ metabolism of *C. autoethanogenum*, the literature lacks a steady-state dataset where carbon flows in a CO_2_+H_2_ fermentation are quantified. Here we aimed to quantify steady-state CO_2_+H_2_ fermentation using fully instrumented chemostats and the model acetogen *C. autoethanogenum*. Subsequently, we showed that CO_2_ is a promising feedstock alternative to CO, as more than half of the substrate CO_2_ carbon was converted into ethanol. Furthermore, supplementation with CO at low concentrations improved fermentation performance significantly.

## Materials and Methods

### Bacterial strain, growth medium, and continuous culture conditions

A derivate of *Clostridium autoethanogenum* DSM 10061 strain—DSM 19630— deposited in the German Collection of Microorganisms and Cell Cultures (DSMZ) was used in all experiments and stored as glycerol stocks at − 80 °C. This non-commercial strain was grown on CO_2_+H_2_ (~23% CO_2_, ~67% H_2_ and ~10% Ar; BOC Australia) and CO/CO_2_/H_2_ (~2% CO, ~23% CO_2_, ~65% H_2_, and ~10% Ar; BOC Australia) in chemically defined medium (Valgepea et al., 2017). Cells were grown under strictly anaerobic conditions at 37 °C and at a pH of 5 (maintained by 5 M NH_4_OH). Chemostat continuous culture achieved steady-states at dilution rates (D) = 0.47 ± 0.01 (CO_2_+H_2_; specific growth rate (μ) = 0.0196 ± 0.0004 [average ± standard deviation]), 0.5 ± 0.01, and 1 ± 0.01 day^−1^ (CO/CO_2_/H_2_; μ = 0.021 ± 0.0004, and 0.042 ± 0.0008 h^−1^ respectively). See Table 1 for steady-state gas-liquid mass transfer rate data. The steady-state results reported here were collected after optical density (OD), gas uptake and production rates had been stable in chemostat mode for at least three working volumes. See Valgepea et al. (2017a) for details on equipment.

**Table 1.**
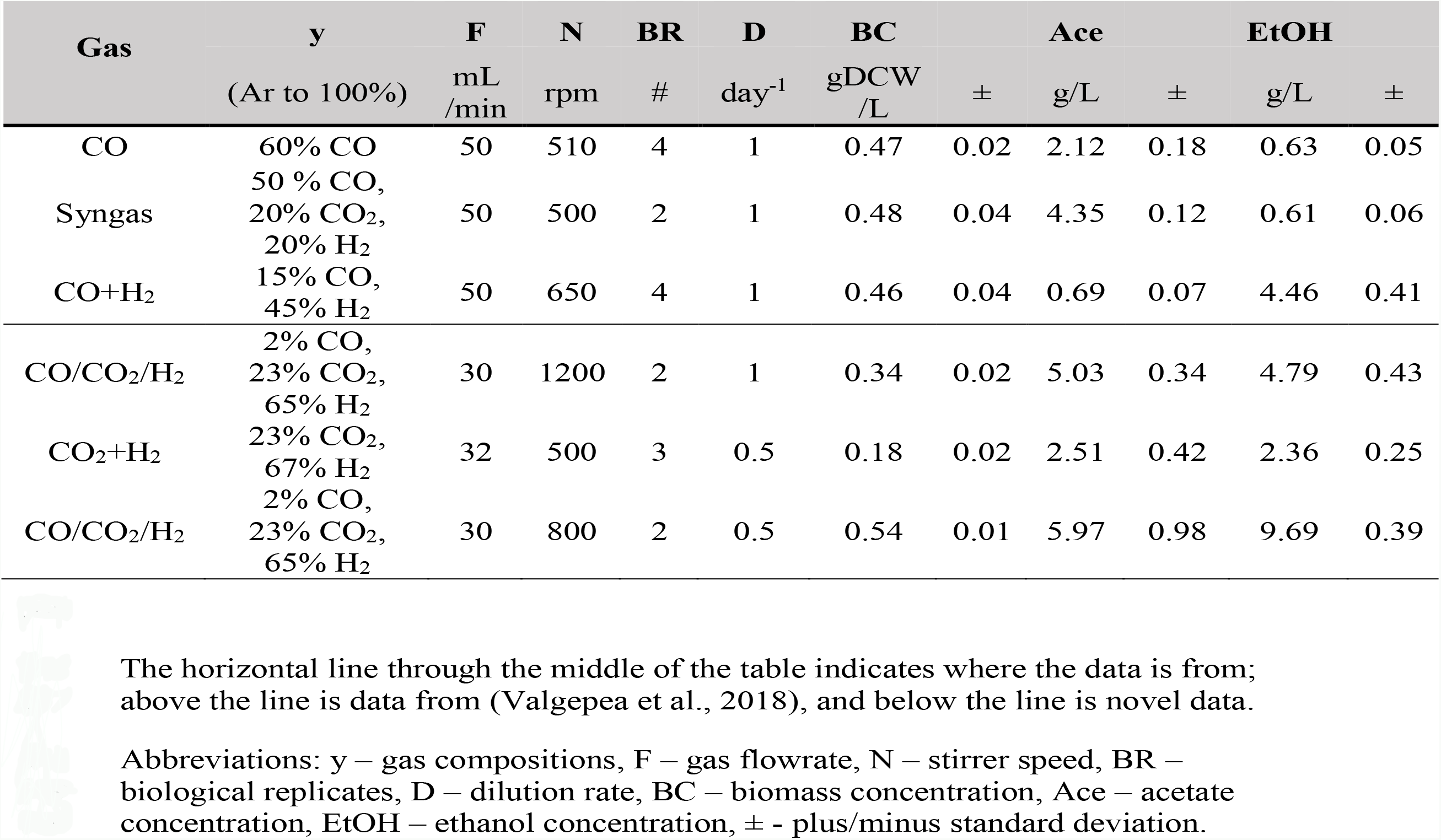
Summary of low-biomass *Clostridium autoethanogenum* fermentations.

## Experimental analysis

### Biomass concentration and extracellular metabolome analyses

Biomass concentration (gDCW/L) was estimated and extracellular metabolome analysis carried out as specified in Valgepea et al. (2018).

### Bioreactor off-gas analysis

Bioreactor off-gas was analyzed by an online Hiden HPR-20-QIC mass spectrometer. The Faraday Cup detector monitored the intensities of H_2_, CO, ethanol, H_2_S, Ar, and CO_2_ at 2, 14, 31, 34, 40, and 44 amu, respectively, in the bioreactor off-gas. These masses were chosen so that each target compound would be represented by a unique signal. This was determined to be essential to achieve the highest confidence in quantification using preliminary experiments as interferences from other compounds at a shared mass could not be reliably accounted for (e.g. the more intense signal from CO at 28 amu could not be used due to the uncertainty of interference at 28 amu from the CO_2_ fragment). Gas from the cylinder was used as the calibration gas for each MS-cycle (i.e. ‘online calibration’) to achieve reliable off-gas analysis (Valgepea et al., 2017). See below for details on quantification of gas uptake and production rates.

## Quantification

### Gas uptake and production rates

Gas uptake (CO, CO_2_ and H_2_) and production (ethanol) were determined using “online calibration” of the MS by analyzing the respective feed gas directly from the cylinder after each analysis cycle of the bioreactors. Specific rates (mmol/gDCW/h) were calculated by taking into account the exact composition of the respective gas, bioreactor liquid working volume, feed gas flow rate, off-gas flow rate (based on the fractional difference of the inert gas [Ar] in the feed and off-gas composition), the molar volume of ideal gas, and the steady-state biomass concentration.

### Carbon balance analysis

The carbon balances were determined at 116 ± 11%, 103 ± 12%, and 108 ± 11% for CO_2_+H_2_, and CO/CO_2_/H_2_ at D = 0.5 and 1 day^−1^ respectively (total C-mol products/total C-mol substrates), as specified in Valgepea et al. (2017).

### Genome-scale metabolic modelling with GEM iCLAU786

Model simulations were performed using genome scale model (GEM) iCLAU786 of *C. autoethanogenum* and flux balance analysis (FBA) (Orth and Palsson, 2011) as specified in Valgepea et al. (2018). Briefly, we used FBA to estimate intracellular fluxes (SIM1–26) and predict “optimal” growth phenotypes for experimental conditions (SIM27–62) using either maximization of ATP dissipation or biomass yield, respectively, as the objective function. Complete simulation results identified as SIMx (e.g. SIM1) in the text are in Supplementary Files. SIM1-19, 27-41, and 49-55 are from Valgepea et al. (2018). In addition to details described in Valgepea et al. (2018), CO_2_ reduction to formate was forced from the formate dehydrogenase (FdhA) reaction scheme (rxn00103_c0, SIM17-30) to the FdhA/Hydrogenase ABCDE complex (HytABCDE) reaction scheme (rxn08518_c0, SIM31-40) when maximizing for biomass formation, as described by Mock et al. (2015). SIM56-62 also stopped export of pyruvate (rxn05469_c0), a decision validated by HPLC data.

## Results

### *Clostridium autoethanogenum* steady-state fermentation of carbon dioxide and hydrogen

*Clostridium autoethanogenum* cells reached steady-state when growing on CO_2_+H_2_ in chemostats at dilution rate (D) ~0.5 day^−1^ (specific growth rate (μ) ~0.02 h^−1^) with a biomass concentration of 0.18 ± 0.02 g dry cell weight (gDCW)/L (Figure 1**A**). It is important to note that attempts to reach a steady-state at D =1 day^−1^ were unsuccessful. Unlike the chemostat cultures of *C. autoethanogenum* with CO (Valgepea et al., 2018; 2017a) and CO_2_+H_2_ retentostat cultures (Mock et al., 2015), the CO_2_+H_2_ cultures could not reach stable biomass concentrations before the culture began oscillation cycles; previously observed above ~1.6 gDCW/L (Valgepea et al., 2017). The physiological reason and mechanism for such oscillatory culture behavior are under investigation, but we assumed that cell recycling is a requirement for CO_2_+H_2_ culture stability. For example, Molitor et al. (2019) showed consistent, high-biomass concentration and high-acetate CO_2_+H_2_ fermentation with *Clostridium ljungdahlii* in a retentostat with complete recycling.

**Figure 1.**
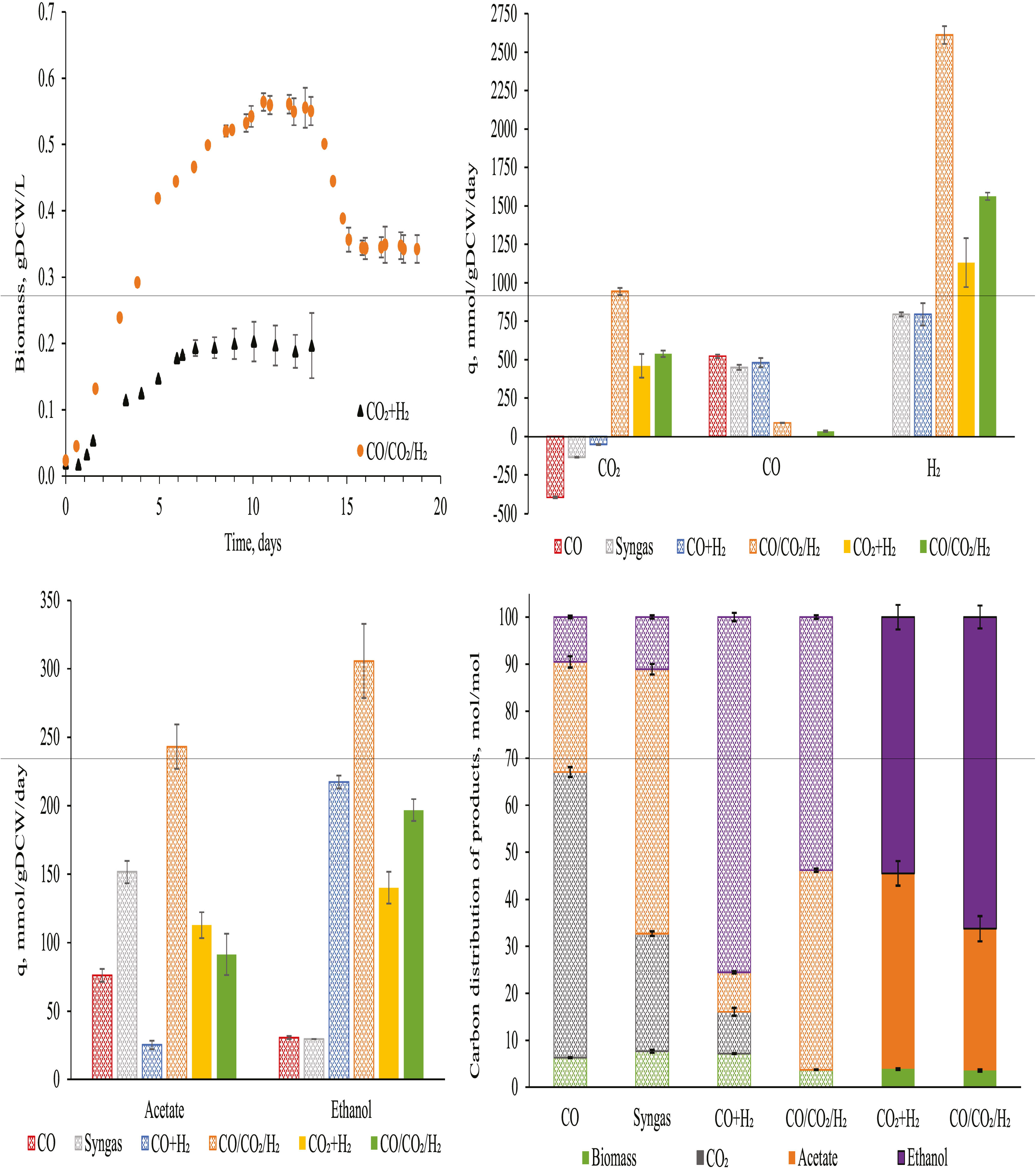
Important fermentation characteristics of *Clostridium autoethanogenum* in autotrophic chemostats. Results from Valgepea et al. (2018) are also displayed (**B**, **C** & **D**), the conditions of all fermentations are summarized in Table 1. Growth curves of novel fermentations with standard deviation at steady-state (**A**). Specific rates of uptake (**B**) and production (**C**) for important metabolites. Product carbon balances (**D**). Values represent the average ± standard deviation between biological replicates. Number of biological replicates, and detailed gas composition for each fermentation are available in Table 1. Patterned bars indicate a D of 1 day^−1^, full bars indicate a D of 0.5 day^−1^ (**B**, **C** & **D**). Abbreviations: *q* ‒specific rate, *DCW* – dry cell weight.

Despite the attempt to reach a steady-state at D = 1 day^−1^, cells reached steady-state at dilution rate = 0.5 day^−1^. Under those conditions, the specific production rates of ethanol and acetate were 140 ± 10 and 113 ± 9 mmol/gDCW/day, respectively (Figure 1**C**). Strikingly, the specific rate of carbon incorporation (i.e. qCO_2_) was 480 ± 80 mmol/gDCW/day (Figure 1**B**), and around half of that carbon was captured as ethanol (54 ± 3 mol%) (Figure 1**D**). Fermentation conditions and titers are available in Table 1, showing an impressive ethanol concentration compared to previous fermentations where CO was the main carbon and energy source.

Despite the different dilution rate, the CO_2_+H_2_ results generated were compared to previously published chemostat cultures of *C. autoethanogenum* grown on CO, syngas, and CO+H_2_ (Valgepea et al., 2018) at similar biomass concentrations (~0.5 gDCW/L) (Figure 1**B**, **C** & **D**). Specific rates of acetate and ethanol production achieved here for CO_2_+H_2_ cultures fell between those for syngas (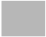) and CO+H_2_ (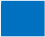) cultures (Figure 1**B** & **D**). However, the specific rate of carbon incorporation was higher for CO_2_+H_2_ (Figure 1**C**). We found that more than half of the captured CO_2_ was converted into ethanol (Figure 1**D**). These results were encouraging, especially as ethanol production has unfavorable stoichiometry compared to acetate (Mock et al., 2015). Furthermore, the H_2_ specific uptake rate (1130 ± 160 mmol/gDCW/day) showed that higher H_2_ uptake rates are achievable (compared to old datasets). These results show that higher carbon yields are possible (Valgepea et al., 2018). To further investigate the metabolic demand and the feasibility of CO_2_+H_2_ fermentation, we utilized the steady-state dataset as constraints for the GEM to find candidate mechanisms for improving CO_2_+H_2_ fermentation using iCLAU786.

### Metabolic model of carbon dioxide and hydrogen fermentation

Estimation of intracellular processes constrained by *in vivo* datasets represents an important developmental step for progressing acetogenic CO_2_ valorization. Here, for instance, comparing CO_2_+H_2_ and CO-containing fermentation fluxes was possible (Figure 2). See Supplementary Files for complete details.

**Figure 2.**
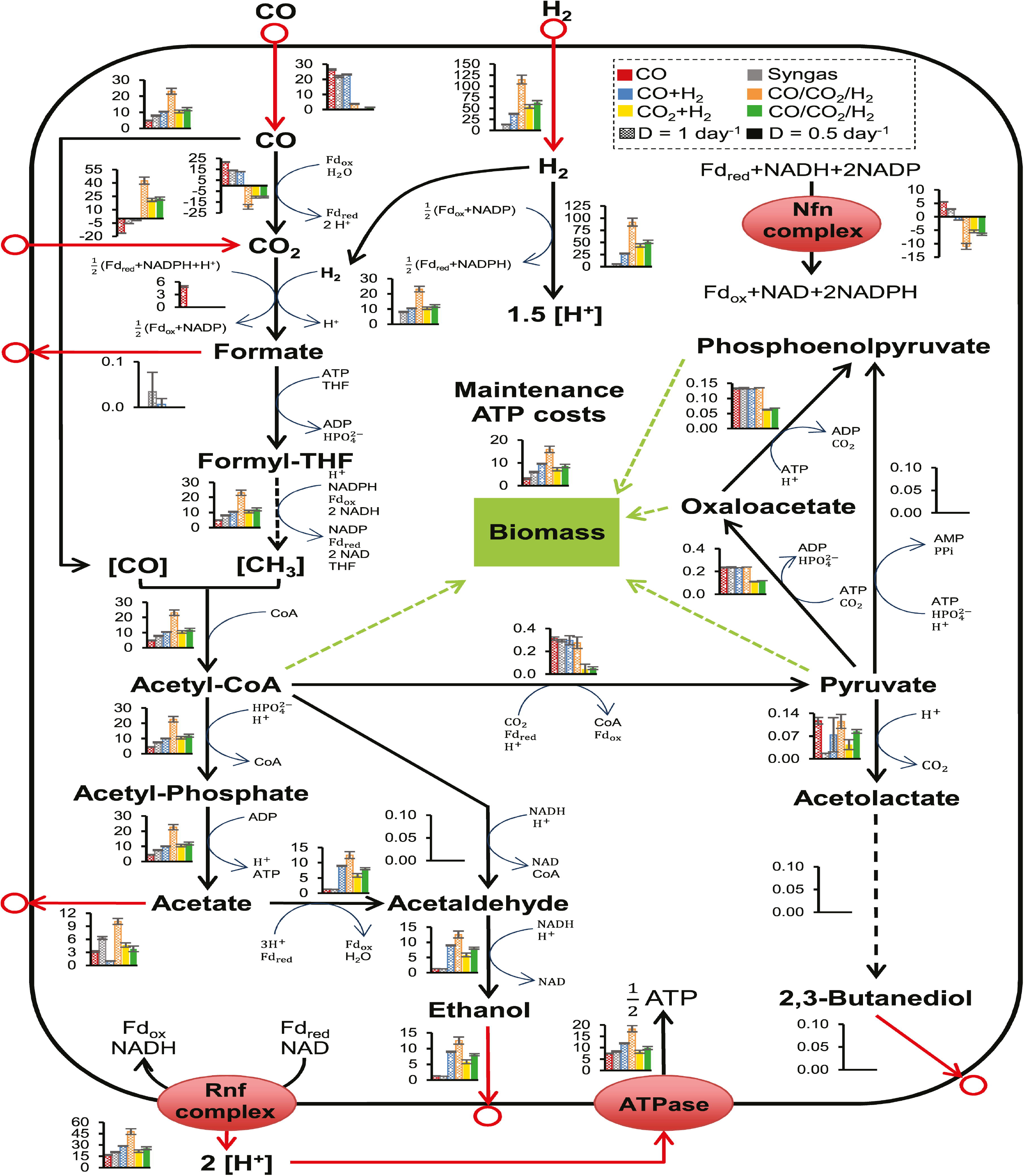
Predictions of central metabolic pathway fluxes for autotrophic fermentations of *Clostridium autoethanogenum* using iCLAU786, flux balance analysis, and chemostat data. Results from Valgepea et al. (2018) are also displayed, the conditions of these fermentations are summarized in Table 1. Fluxes (mmol/gDCW/h) are represented as the average ± standard deviation between biological replicates. Number of biological replicates, and detailed gas composition for each fermentation are available in Table 1. Arrows show the direction of calculated fluxes; red arrows denote uptake or secretion, dashed arrows denote a series of reactions. Brackets denote metabolites bound by an enzyme. Refer to Supplementary Files for enzyme involvement, metabolite abbreviations, and complete flux balance analysis datasets.

Intracellular metabolite fluxes from the FBA showed remarkable similarity to the combined theoretical stoichiometry of acetate and ethanol production (Mock et al., 2015) and indicated energetic cofactor circuits with mapping close to 1:1 (experimental:theoretical stoichiometry; Supplementary Files). Ethanol production likely occurred *via* acetaldehyde:ferredoxin oxidoreductase (AOR; leq000004) under autotrophic conditions, with the HytABCDE (leq000001) and Nfn complex (leq000002) likely facilitating cofactor production *via* electron bifurcation (Figure 2) (Valgepea et al., 2018). This is a mechanism for minimization of free energy loss employed by *C. autoethanogenum* and may play a key role in sustaining proton motive force by balancing acetate, ethanol, and ATP production (Mock et al., 2015; Valgepea et al., 2018). Engineering acetogens to redirect this energy towards cellular growth, sacrificing some ethanol production, could be beneficial for CO_2_ fermentation (Emerson and Stephanopoulos, 2019).

It was notable that, unlike CO fermentations, the pyruvate:ferredoxin oxidoreductase (PFOR; rxn05938_c0; acetyl-CoA ↔ pyruvate) flux was not significantly in the direction of pyruvate (Figure 2) (Valgepea et al., 2018). Under autotrophic conditions, PFOR links the WLP to anabolic pathways associated with biomass (Furdui and Ragsdale, 2000), and therefore this indicated high cell-specific energetic limitation.From this observation, we hypothesized that CO supplementation could provide a potential solution, as CO oxidation would generate Fd_red_. Furthermore, an ATP/H_2_ flux ratio of ~0.15 was observed here compared to an ATP/CO ratio of ~0.28 in CO only fermentations (Valgepea et al., 2018). Considering CO+H_2_ and CO_2_+H_2_ fermentations had equal carbon-flux through the WLP (~10 mmol/gDCW/h; Figure 2), supplementation with renewable CO from CO_2_ electrolysis could control biomass formation and culture stability. A similar process (but CO fermentation) was detailed by Haas et al. (2018).

### *Clostridium autoethanogenum* steady-state fermentation of carbon dioxide and hydrogen supplemented with carbon monoxide

To validate our modelling hypothesis, *Clostridium autoethanogenum* was cultured with a low concentration of carbon monoxide in addition to CO_2_ and H_2_ (CO/CO_2_/H_2_) in chemostats. A steady-state was reached at D = 0.5 day^−1^ (μ ~0.02 h^−1^), and at D =1 day^−1^ (μ ~0.04 h^−1^; Figure 1**A**; biomass concentrations of 0.54 ± 0.01 and 0.34 ± 0.02 gDCW/L respectively). CO/CO_2_/H_2_ fermentations at a D = 1 day^−1^ (CO/CO_2_/H_2_^1^) and a D = 0.5 day^−1^ (CO/CO_2_/H_2_^0.5^) showed simultaneous uptake of CO (89 ± 2 and 36 ± 4 mmol/gDCW/day, respectively) and CO_2_ (940 ± 20 and 540 ± 20 mmol/gDCW/day, respectively) (Figure 1**B**). The co-utilization of both C1 gases is, to the best of our knowledge, an unquantified phenomenon. This led to a specific carbon incorporation (CO/CO_2_/H_2_^1^ – 1030 ± 30 mmol/gDCW/day) larger than any other gas type (maximum of ~450 mmol/gDCW/day for fermentations with CO in Valgepea et al. (2018) or CO_2_+H_2_ in this work). This also resulted in significant improvements to culture performance compared to CO_2_+H_2_ fermentations.

Compared to CO_2_+H_2_, CO/CO_2_/H_2_^0.5^ showed higher acetate and ethanol titers (Table 1) and specific productivities (Figure 1**C**), and a higher ethanol/acetate ratio (2.15 vs 1.24 mol/mol respectively). While at a similar biomass concentration (CO/CO_2_/H_2_^1^ best comparison due to similarity in dilution rate), acetate and ethanol titers (Table 1), and specific productivities (Figure 1**C**) are greater than during fermentation of other CO-containing gases. When comparing to high biomass (~1.4 gDCW/L) CO cultures, CO-supplementation still performs impressively – CO+H_2_ fermentation achieved a higher ethanol titer (11.6 ± 0.4 g/L), while CO and syngas fermentations were similar (3.9 ± 0.2 and 5.4 ± 0.3 g/L respectively). Otherwise, all specific productivities were higher for CO/CO_2_/H_2_^1^ (Supplementary Files). Furthermore, the distribution of carbon to ethanol was still greater than 50% (Figure 1**D**; 53.8 ± 0.4 % and 66 ± 2% for CO/CO_2_/H_2_^1^ and CO/CO_2_/H_2_^0.5^ respectively).

To understand the metabolic effects of supplementing CO, FBA was performed using the same conditions and alterations as for CO_2_+H_2_ (Figure 2). Notably, the WLP specific flux throughput for CO/CO_2_/H_2_^1^ was ~2-fold greater than for any other gas type (including high-biomass [Valgepea et al., 2018]). Furthermore, for CO_2_ fermentations, Nfn complex flux direction was opposite that of CO and syngas fermentations. CO/CO_2_/H_2_^0.5^ also showed significantly greater flux through the AOR, whilst specific WLP productivity was insignificantly different compared to CO_2_+H_2_.

## Discussion

Achieving steady-state continuous cultures using CO_2_+H_2_ mixtures, without cell recycling here, was challenging. Yet, compared to other organisms fermenting CO_2_+H_2_ with continuous medium exchange, *Clostridium autoethanogenum* performs well (Table 2). No direct comparisons can be made to other experiments due to variations in conditions, but *C. autoethanogenum* clearly achieves the highest ethanol production, with comparable quantities of carbonous products also. *Acetobacterium woodii*, along with *Sporomusa ovata*, were shown to perform well when compared to a wide range of acetogens under batch CO_2_+H_2_ conditions (Groher and Weuster-Botz, 2016). Yet, as evidenced by omission of *S. ovata* from Table 2, few continuous culture characterizations of acetogens are available – an essential step for validation of industrial robustness in gas fermentation. As discussed by Molitor et al. (2019), the lack of yeast extract or C_≥2_ substrates is also distinguishing between fermentations.

**Table 2.**
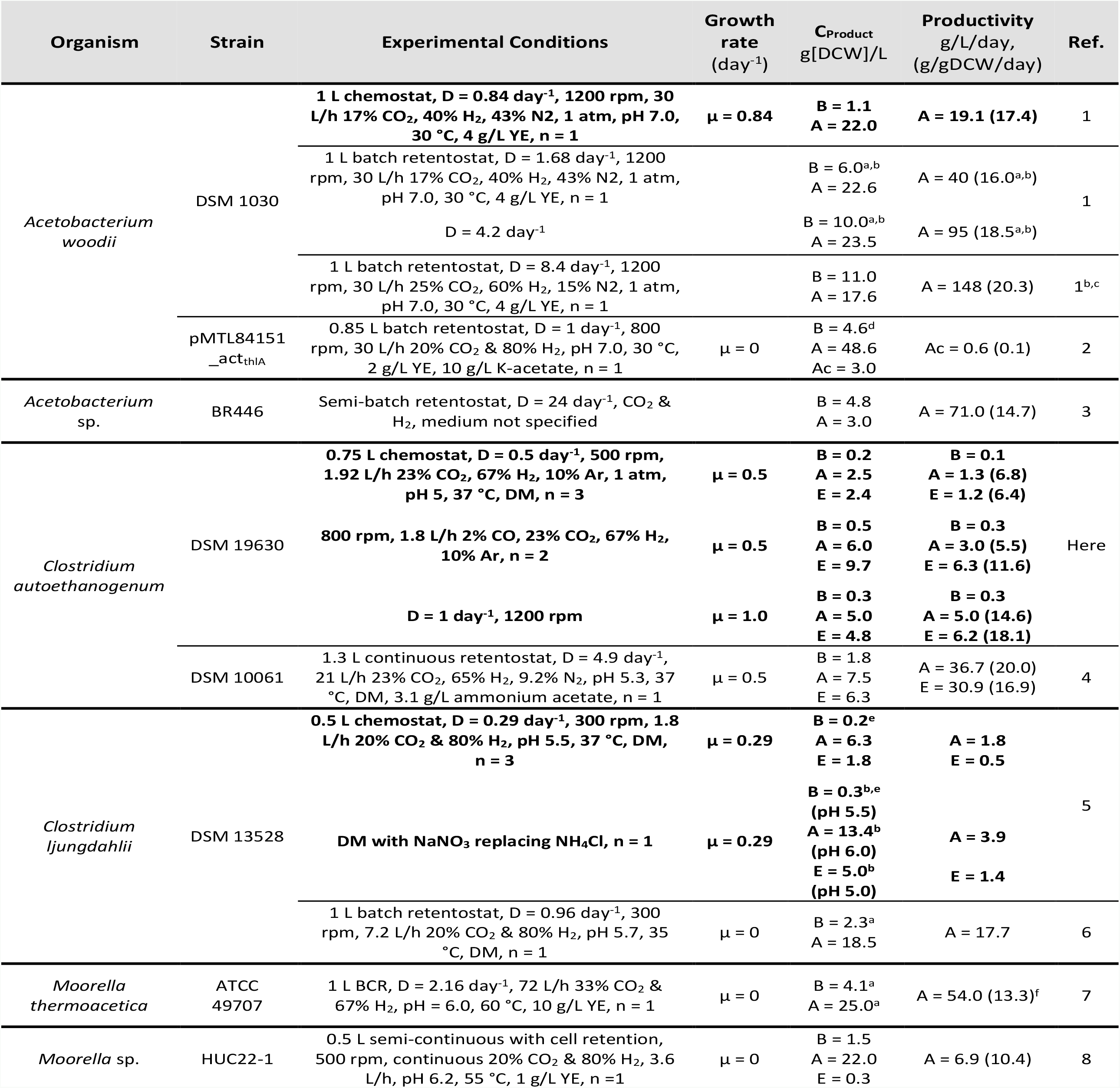
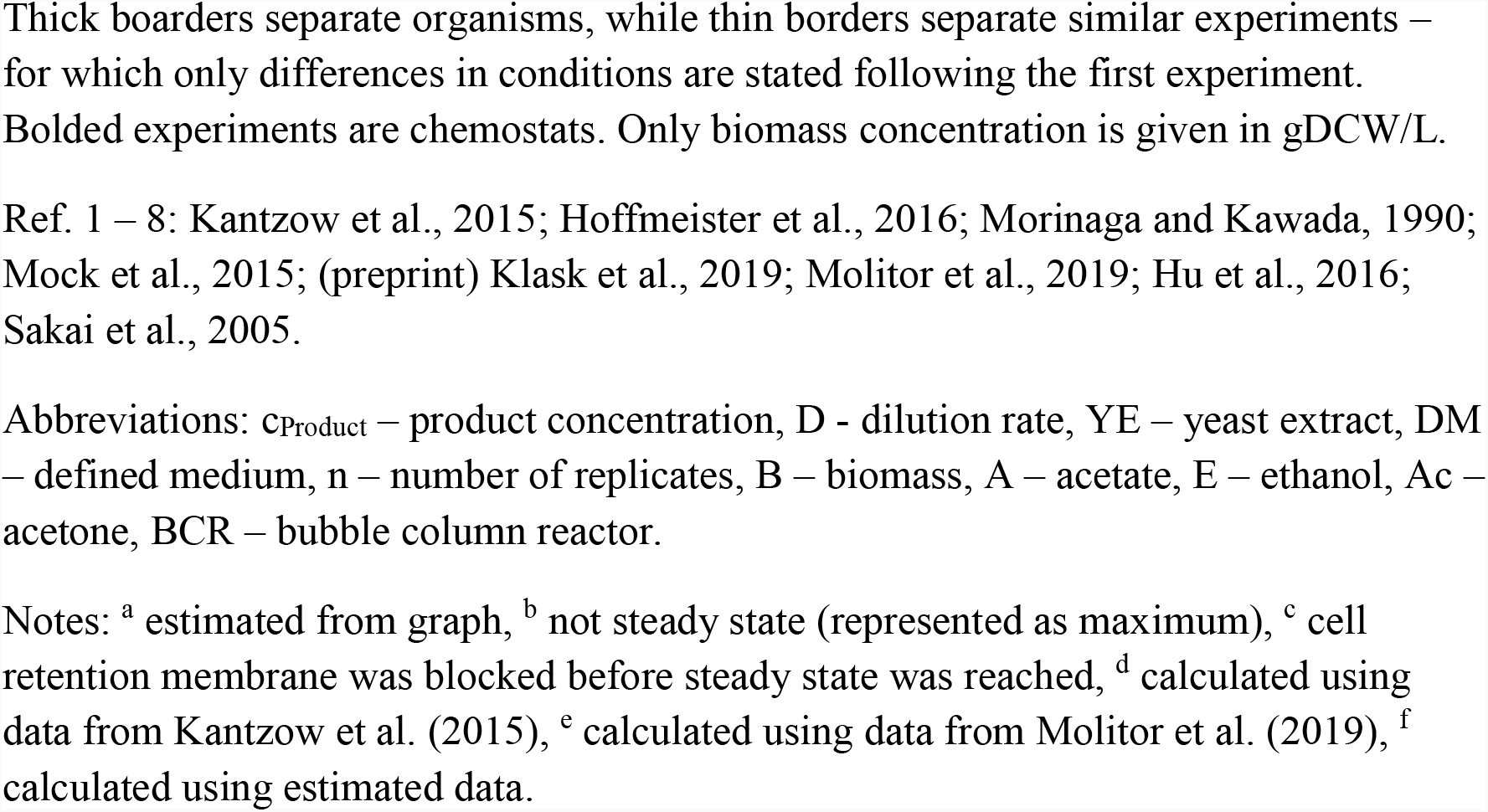
Summary of quantitative and continuous CO_2_+H_2_ fermentations.

Notably, CO_2_+H_2_ cultures displayed higher variability between biological replicates compared to those of CO-containing gas mixtures (Figure 1) (Valgepea et al., 2017). This may indicate variable organism fitness, a trait previously discussed for *C. autoethanogenum* by Liew et al. (2016), who extensively covered numerous techniques used for enhancing gas fermentation including – coupling to other processes, adaptive laboratory evolution, and metabolic engineering of acetogens using genetic tools. CO-supplementation could be a valuable option for enhancement as it overcomes inherent problems linked to engineering acetogens. Supplementation of low quantities of CO here stabilized the culture, enabled culturing at D =1 day^−1^, and achieved higher biomass concentration with a carbon incorporation larger than any other gas type – all without compromising by-product distribution.

While Valgepea et al. (2018) found that syngas fermentation lead to CO-only fermentation at steady-state, we observed co-utilization of CO and CO_2_. Tizard and Sechrist (2015) have also shown co-utilization for *C. autoethanogenum* continuous cultures, and it seems that co-uptake may also occur for some points of syngas batch fermentation ([preprint] Infantes et al., 2020). Co-utilization of sugars was found for *E. coli* in chemostats – where inhibition of consumption, but no change in induction time was observed (Standing et al., 1972). The WLP is most likely no different, in that metabolism of CO is preferential, yet the pathway can co-consume CO_2_ under certain conditions.

Various efforts have been made towards enhancing CO_2_(+H_2_) fermentation to C_≥2_ products (Table 2) (Emerson and Stephanopoulos, 2019). Braun and Gottschalk (1981) first discovered the potential for enhancement when *Acetobacterium woodii* simultaneously consumed fructose and a headspace of CO_2_+H_2_ during batch cultivation. Growth and acetate production was high but no characterization of the headspace was performed. More recently, continuous glucose-supplemented CO_2_+H_2_ fermentation of *Moorella thermoacetica* by Park et al. (2019) did not lead to net uptake of CO_2_. Furthermore, Jones et al. (2016) did not show net CO_2_ uptake for a wide range of acetogens (not *A. woodii*) fermenting syngas and fructose. *A. woodii* generates a sodium ion (Na^+^) gradient (Hess et al., 2013) rather than a proton (H^+^) gradient for membranous ATP generation (Bengelsdorf et al., 2018; Pierce et al., 2008; Poehlein et al., 2015). This may highlight an important metabolic difference from other model acetogens – decoupling the resources of the WLP and membranous ATP generation pathways could facilitate fermentation of sugar and CO_2_+H_2_ simultaneously.

Other enhancements have also struggled to achieve net CO_2_ uptake. Co-culture of *C. acetobutylicum* and *C. ljungdahlii* showed syntrophic metabolic coupling when fermenting glucose, fructose, and CO_2_+H_2_, but no net CO_2_ uptake (Charubin and Papoutsakis, 2019). Addition of nitrate to batch CO_2_+H_2_ fermentation by *C. ljungdahlii*, increased biomass concentration and subsequently volumetric productivity of acetate (Emerson et al., 2019). However, the specific WLP productivity decreased, meaning lower utilization of CO_2_. Other organisms not recognized as gas fermenters can also use mixotrophy to minimize carbon loss, such as *Clostridium beijerinckii* but have not displayed net CO_2_ uptake either (Sandoval-Espinola et al., 2017). To the best of our knowledge, this is the first report where supplementation of a substrate other than H_2_, increased productivities of continuous acetogenic CO_2_ fermentation while maintaining net CO_2_ utilization. Furthermore, the effect of CO supplementation on CO_2_ utilization was superlinear, indicating a synergistic mechanism (Park et al., 2019). This is encouraging for development of bioprocesses valorizing CO_2_.

Comparisons between fermentation datasets enables us to speculate about the positive effect of CO-supplementation on CO_2_+H_2_ fermentation. Although, addition of CO led to minimal metabolic shifts (Figure 2 – CO_2_+H_2_ vs CO/CO_2_/H_2_^0.5^ and Supplementary Files), FBA showed that CO supplementation caused significant increases to the reduced ferredoxin consumption by AOR and Rnf complex (leq000004 and M002, respectively) compared to CO_2_+H_2_ (Figure 2). The overflow model proposed by Richter et al. (2016) suggests that high NADH production *via* Rnf and Nfn complexes (leq000002) is also important for reducing AOR product inhibition. In this way, NADH facilitates fast metabolism of acetaldehyde to ethanol *via* alcohol dehydrogenase (Adh(E); rxn00543_c0). Decreasing the acetate concentration reduces acidification and the ATP cost for excreting acetate (Valgepea et al., 2018). Including acetaldehyde conversion to ethanol and association to acetic acid, this also leads to consumption of 2 H^+^ (4 here vs 2 produced *via* CODH). Therefore, CO consumption decreases the intracellular H^+^ pool, and following Le Chatelier’s principle, drives HytABCDE activity. Indeed, the change in specific H_2_ uptake relative to specific CO_2_ uptake is greater than that of CO (for CO_2_+H_2_ vs CO/CO_2_/H_2_ at D = 0.5 day^−1^, Supplementary Files). Subsequently, the relative gain in free energy from H_2_ is ~ 4-fold greater than CO. We speculate this is ultimately responsible for the improved fitness of CO-supplemented CO_2_+H_2_ fermentation by *C. autoethanogenum*. We propose the following five critical factors to this enhanced metabolism: [1] metabolism of CO increases the intracellular pool of reduced ferredoxin; [2] this stimulates oxidation of ferredoxin, which if consumed by the AOR; [3] reduces ATP costs; and [4] decreases the H^+^ pool/acidification; which therefore [5] drives H_2_ uptake for further reduction of ferredoxin. Evidently, additional understanding of acetogenic redox metabolism, from a thermodynamic perspective, is important for developing acetogenic CO_2_-valorization as a platform industrial bioprocess (Cueto-Rojas et al., 2015).

Physicochemical properties could also play a key role in CO-supplementation enabling to achieve a stable CO_2_+H_2_ chemostat culture at D =1 day^−1^. Generation of a stable and large non-equilibrium is what drives microbial growth (Igamberdiev and Kleczkowski, 2009; Qian and Beard, 2005; Quéméner and Bouchez, 2014) and gas-liquid mass transfer (Ma et al., 2005). For continuous culture of gas fermenting microbes, an inherent relationship between substrate mass transfer and culture growth exists (Supplementary Files). An important parameter for these systems is the Gibb’s free energy of a system (Cueto-Rojas et al., 2015). This describes the thermodynamic favorability of the reaction system – termed spontaneity. Here, analysis of experimental flux and Gibbs free energy suggests that CO_2_+H_2_ fermentation is infeasible 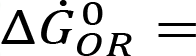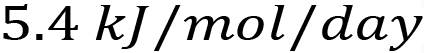, whereas CO-supplemented CO_2_+H_2_ fermentation is feasible 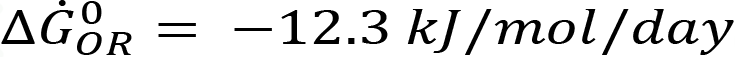; Supplementary Files). Though these calculations use standard conditions, they do indicate how close CO_2_+H_2_ fermentation is to the thermodynamic limit of metabolism. Theoretically, minute and unobservable changes to chemostat CO_2_+H_2_ fermentation can disrupt the culture (Henry and Martin, 2016). Thus, increasing the free energy of central metabolism with CO-supplementation appears to keep metabolism in a spontaneous and stable state by increasing reduced ferredoxin production.

The mechanisms for achieving the 2-fold higher specific WLP flux throughput for CO/ CO_2_/H_2_^1^ compared to others is less clear but appears to be linked to the difference in primary substrate. CO/CO_2_/H_2_^1^ and CO+H_2_ are the most similar CO_2_ and CO fermentations, respectively (D ~1 day^−1^ and carbon to hydrogen feed ratio (~1:3); Table 1), and the maximum carbon incorporation per cell for CO+H_2_ was roughly half of that of CO/CO_2_/H_2_^1^ (~450 vs ~1000 mmol/gDCW). Theoretically, cells will maximize carbon-to-redox metabolism by minimizing thermodynamic losses. CO supplementation to a CO_2_+H_2_ culture seems to facilitate this as (H_2_/carbon)_feed_ − (H_2_/carbon)_flux_ was ~0 mol/mol for CO/CO_2_/H_2_ fermentations only (Supplementary Files) – an indication of the relative magnitude of carbon and redox metabolism. This suggests that high specific fluxes for CO/CO_2_/H_2_^1^ may be a result of (close to) optimal co-factor recycling by *C. autoethanogenum*’s WLP and redox pathway. Thus, the lower energy associated with CO_2_ fermentation may, counterintuitively, stimulate specific WLP activity when in the presence of appropriate energy-containing substrates. Further quantifications of CO_2_ metabolism and characterizations of enzyme activities are required to confirm these hypotheses (Supplementary Files), as they assist our ability to engineer the links between redox and carbon metabolisms.

We established a dataset quantifying steady-state of the model acetogen *C. autoethanogenum* during autotrophic-CO_2_+H_2_ growth in chemostat cultures. This enabled analysis *via* FBA, and highlighted CO as a potential supplement. CO supplementation successfully improved metabolic stability and CO_2_ utilization. This was the first time that intracellular fluxes for net uptake of CO_2_ (with enhancement) where characterized. Industry is actively developing gas fermentation to valorize CO_2_ (Haas et al., 2018 & Tizard and Sechrist, 2015). Previously, genetic and process engineering of gas fermentation successfully developed the technology for industrial CO valorization (Liew et al., 2016). Therefore, progression to industrial CO_2_ valorization is foreseeable, and CO supplementation may play a role in the continuing diversification of industrial gas fermentation.

## Author Contributions

All authors viewed and approved the manuscript. All authors contributed significantly to the work. KV, EM, and LN conceived the project. JH, KV and EM designed the experiments and analysed the results. JH and KV performed experiments, supported by RL, IC, MP, and EM. JH wrote the manuscript with the help of KV, EM, RT, SS, MK, and LN.

## Acknowledgments

Elements of this research utilized equipment and support provided by the Queensland node of Metabolomics Australia, an initiative of the Australian Government being conducted as part of the NCRIS National Research Infrastructure for Australia. IC would like to acknowledge support from the German Academic Exchange Service (DAAD) through the “DAAD Kurzstipendien für Doktoranden”. We thank the following investors in LanzaTech’s technology: Sir Stephen Tindall, Khosla Ventures, Qiming Venture Partners, Softbank China, the Malaysian Life Sciences Capital Fund, Mitsui, Primetals, CICC Growth Capital Fund I, L.P. and the New Zealand Superannuation Fund. There was no funding support from the European Union for the experimental part of the study. However, KV acknowledges support also from the European Union’s Horizon 2020 research and innovation programme under grant agreement N810755.

## Contribution to the field

Acetogenic bacteria comprise an ancient lineage and play a major role in global carbon cycle (accounting for at least 10^13^ kg of acetate produced annually and 20% of the fixed carbon on earth). Due to their ability to grow autotrophically on carbonous waste-gas feedstocks, these organisms have gained significant interest in biotechnological applications. However, acetogens are considered living at the thermodynamic edge of life when growing autotrophically. Although they have evolved sophisticated strategies to conserve energy from reduction potential differences between major redox couples, this coupling is sensitive to small changes in thermodynamic equilibria. In the manuscript, we present experimental data showing CO_2_ conversion to ethanol by an acetogenic bacteria used for industrial scale gas fermentation. Furthermore, we showed that supplementing CO enhances CO_2_+H_2_ fermentation performance significantly. Analysis was only possible due to the first rigorously quantified dataset from continuous CO_2_ and H_2_ fermentation. This enabled discovery of notable insights into metabolic function – providing a potential guide for metabolic engineering. Therefore, here we outline that *Clostridium autoethanogenum* offers a promising route for the sustainable production of fuels and chemicals from a wide range of waste feedstocks – including CO_2_.

## Supplementary Files

The Supplementary Files for this article can be found online.

