## Supplementary Materials and Methods, Tables S1-9, and Figures S1-4 for "Enhancing CO_2_-valorization using *Clostridium autoethanogenum* for sustainable fuel and chemicals production"

### Data analysis

#### Analysis of flux data

Analysis of flux data from flux balance analysis (FBA) was performed to assist hypothesis formation. In relation to Table S7, flux data was normalized by $CO_2$ uptake (rxn05467_c0) before the differential change was calculated. Column headings indicate what flux was subtracted from one another. In relation to Table S8 & S9, and Figure S2, flux data was normalized by carbon uptake (i.e. CO uptake [rxn10480_c0] plus $CO_2$ uptake [rxn05467_c0]) $CO_2$ flux was included only if an uptake was present. Thus, for fermentations primarily fermenting CO, production of $CO_2$ from CO was not subtracted from carbon uptake. Relative fluxes were calculated by dividing flux $ij$ by the respective (normalized) flux of $CO/CO_2/H_2$[1] – where $i$ represents any fermentation dataset except for $CO/CO_2/H_2$[1], and $j$ represents any reaction. Similarly, differential fluxes were calculated by taking flux $ij$ and subtracting the respective (normalized) flux of $CO/CO_2/H_2$[1].

#### Gibb's free energy balance

A Gibb's free energy balance was conducted based on Liu et al. (2016). Free energy reactions from $CO_2$ to product ($\Delta G^0_{CO_2 \rightarrow product}$) were used to energetically describe each reaction. $CO_2$ was chosen as the base carbon molecule, as CO can readily be converted to $CO_2$ by cells. Volumetric free energy flow rates were calculated rather than amount of energy (i.e. kJ/L/day vs. J). Therefore, the $\Delta G^0_{CO_2 \rightarrow product}$ for each reaction was multiplied by its corresponding volumetric molar flow rate ($\dot{n}_{product}$ [mol/L/day]) from Table S6, to obtain a change in Gibb's free energy flowrate for the overall system ($\Delta \dot{G}^0_{OR}$[kJ/L/day]). A biomass molecular weight of 24 g/mol was used to obtain $\dot{n}_{biomass}$ (Valgepea et al., 2017). $\Delta_r G^0_{CO_2 \rightarrow product}$ for biomass formation [1] was taken from Liu et al. (2016), while ethanol [2] and acetate formation [3] were found by reversing their respective combustion reactions, and combustion of $H_2$ [4] and CO [5] were used for their respective reactions. Aside from biomass, Chemistry Reference contained the appropriate $\Delta G^0_R$ information.

$$CO_2 + 0.24\,NH_3 + 0.525\,H_2O \rightleftharpoons CH_{1.77}O_{0.49}N_{0.24} + 1.02\,O_2 \qquad [1]$$
$$\Delta G^0_{\mathrm{R}} = 479\,kJ/mol$$

$$2\,CO_2 + 3\,H_2O \rightleftharpoons CH_3CH_2OH + 3\,O_2 \qquad [2]$$
$$\Delta G^0_{\mathrm{R}} = 1300\,kJ/mol$$

$$2\,CO_2 + 2\,H_2O \rightleftharpoons CH_3COOH + 2\,O_2 \qquad [3]$$
$$\Delta G^0_{\mathrm{R}} = 873\,kJ/mol$$

$$H_2 + {}^1\!/_2\,O_2 \rightleftharpoons \mathrm{H}_2O \qquad [4]$$
$$\Delta G^0_{\mathrm{R}} = -230\,kJ/mol$$

$$CO + {}^1\!/_2\,O_2 \rightleftharpoons CO_2 \qquad [5]$$
$$\Delta G^0_{\mathrm{R}} = -257\,kJ/mol$$

Typically, concentrations, not fluxes, are used for these calculations. However, absolute quantification of some intracellular metabolite concentrations are difficult to obtain (Lange et al., 2001 & Taymaz-Nikerel et al., 2009). To say definitively whether a reaction is spontaneous or not ($-\Delta G^0_R$ and $\Delta G^0_R$ respectively), one would have to know the intracellular concentrations. 0 kJ/mol is therefore the switch point between a reaction being spontaneous and non-spontaneous. Importantly, time only affects the magnitude of the value, as it must be positive – thus, the rule holds when using rate values (it may be important to compare similar dilution rates though). We know that the system is negative $\Delta G$ because we see growth and/or conversion of substrates to products. Here, the standard conditions are assumed, meaning the calculations are not quantitative absolutely, but between comparable datasets results are quantitative relatively.

#### Titer conversion to specific production rate and retention factor correction

Using methods from Villadsen et al. (2011) (pg. 401), the titers from Mock et al. (2015) were converted to chemostat-equivalent specific production rates. Titers were 1.83, 6.3 and 7.48 g/L for biomass, ethanol and acetate respectively. Gas stripping of ethanol was found to be only 1% of total ethanol production here, therefore having minimal impact on the results reported for retentostat culture (not analyzed by Mock et al. [2015], based on $CO/CO_2/H_2$ cultures due to higher ethanol titers). Dilution rates were 0.5 and 4.9 day$^{-1}$ for biomass and metabolites respectively. These were assumed to be growth rate ($\mu$) and dilution rate ($D$), respectively, in the equation $\mu = D(1 - \Omega)$, where $\Omega = 0.9$ was the retention factor. It was assumed that the metabolites from the bleed stream were retained, leading to a productivity increase factor of $(1 - \Omega)^{-1}$. E.g. for ethanol the specific production rate was calculated by;

$$q_{EtOH} = (1 - \Omega)\frac{D \times n_{EtOH}}{x} = 37.4\, mmol/gDCW/day$$

Where $n_{EtOH}$ was the molar concentration of ethanol, $x$ was the biomass concentration of the reactor, and $q$ was the specific production rate of ethanol. When converted, Mock et al. (2015) observed lower specific productivities for retentostat $CO_2$+$H_2$ cultures of *C. autoethanogenum* (DSM 10061) compared to our chemostat fermentations with DSM 19630 (140 ± 10 vs 37, and 113 ± 9 vs 34 mmol/gDCW/day respectively), while the ethanol/acetate ratio was similar (1.2 vs 1.1 mol/mol respectively). However, the advantage of retentostat culture is volumetric production, hence their frequent use in $CO_2$ fermentation (Table 2). However, Mock et al.'s study focused on the biochemical reactions of gas fermentation, and unfortunately, a mass balance was not attempted. Thus, meaningful metabolic comparisons between $CO_2$+$H_2$ chemostats and retentostats are not possible yet; as both the strain of *C. autoethanogenum* and the culture conditions are different between Mock et al. and the data reported here.

### Notes

#### Growth feedback loop and the mass transfer rate-dilution rate relationship

Non-equilibrium thermodynamics have been theorized to dictate metabolism and cell growth (Igamberdiev and Kleczkowski, 2009; Qian and Beard, 2005; Quéméner and Bouchez, 2014). Gas fermenting microbes create a mass transfer, non-equilibrium feedback loop, in that increasing biomass proportionally increases gas consumption – generating further increases in the mass transfer non-equilibrium driving force (Ma et al., 2005). Thus, continuous culture steady-states of gas fermenting microbes exemplify the principle of stable, non-equilibrium thermodynamics. Microbial growth intrinsically links mass transfer rate (MTR) and dilution rate (D) – design parameters known to be important for optimization of continuous gas fermentation performance (Klasson et al., 1991 & 1992). Furthermore, we have now explicitly shown that changes to MTRs (Valgepea et al., 2017a & 2018), and now D (Figure 2), lead to metabolic shifts. Substrate partial pressure, bubble size, and contact time are variables that are controllable and affect MTRs significantly (Bouaifi et al., 2001; Chen et al., 2018; Li et al., 2019). Changes to stirring speed and partial pressure caused a MTR difference between $CO_2$+$H_2$ and $CO/CO_2/H_2^{0.5}$. A ~5-fold increase in ethanol volumetric productivity was observed for $CO/CO_2/H_2^{0.5}$ compared to $CO_2$+$H_2$, while biomass concentration and acetate volumetric productivity observed a ~2-fold increase (Table 2). Yet, the specific flux through the WLP was not significantly affected, agreeing with results from Valgepea et al. (2017a). Varying results have been obtained with elevated partial pressures of $H_2$ and $CO_2$ (Table 2) (Demler and Weuster-Botz, 2011; Kantzow and Weuster-Botz, 2016; Oswald et al., 2018). However, the $H_2$:$CO_2$ ratios used in these studies were 2:1, therefore potentially leading to an $H_2$-limited fermentation. Here and by others (Table 2), $H_2$ to $CO_2$ consumption can be greater than 2:1. Thus, the hypothesis of bicarbonate acidification by Oswald et al. (2018) (also seen by Kantzow and Weuster-Botz [2016]), may only hold at this substrate feed ratio. This highlights an important difference between CO and $CO_2$ fermentations – the necessity of an energy substrate (for $CO_2$) creates an imperative balance between carbon and redox metabolisms. Here, there was no observable improvements to MTRs compared to Valgepea et al. (2018) (shown by volumetric flux in Table S6). To improve MTRs outside of optimizing the MTR-D relationship or without decreasing specific productivities, it may be necessary to reduce any metabolic limitations (i.e. product inhibition) through methods like synergistic co-culture (Charubin and Papoutsakis, 2019; Richter et al., 2016a).

Increases in dilution rate and MTR led to the $CO/CO_2/H_2^1$ steady-state, indicating (when considering $CO_2$+$H_2$ vs $CO/CO_2/H_2^{0.5}$) that specific growth rate (result of D) was most likely the cause of the WLP specific productivity increase. Previously, different dilution rates were not reported for *C. autoethanogenum* chemostats, and there appears to be a proportional relationship between the WLP flux and D. The ethanol/acetate ratio of $CO/CO_2/H_2^1$ also decreased by close to half compared to $CO/CO_2/H_2^{0.5}$, and approached that of $CO_2$+$H_2$ fermentation (Figure 1). This suggests that metabolism was approaching a metabolic state similar $CO_2$+$H_2$ fermentation, and once again may be struggling to balance redox. One solution to this could be increasing CO supplementation to facilitate higher biomass and ethanol production (as seen for $CO_2$+$H_2$ vs $CO/CO_2/H_2^{0.5}$). Additionally, retentostat or multi-stage chemostat set-ups can reduce growth rate pressure while also reducing product inhibition (Table 2) (Mock et al., 2015; Molitor et al., 2019; Richter et al., 2013). This tends to increase the volumetric-, but reduce the specific-productivity, a trade-off that is generally desirable industrially. Furthermore, CO supplementation could increase volumetric productivities for $CO_2$+$H_2$ retentostats. However, the feasibility of attaining the same fold-increase in retentostat cultures is yet to be seen.Thus, further work towards optimization of these parameters may enhance not only the reactor performance, but also the metabolic performance.

### Enzymes characterizations of interest

*C. autoethanogenum* possesses many vitally important enzymes. However, within the acetogen community, there is a diversity of these enzymes, and even more so for some present in the wider communities. We briefly discussed the Rnf complex diversity within acetogens, with more in-depth analysis by Bengelsdorf et al. (2018) and Poehlein et al. (2015). The following discusses other enzymes with notable importance and potential for further characterization.

#### CODH

Enzyme specific activity may play a role in the WLP increase. Mock et al. (2015) found that between CO and $CO_2$+$H_2$ cultures, the largest (~2-fold) significant activity increase was in carbon monoxide dehydrogenase (CODH). Clearly, CODH plays an important role in $CO_2$ fermentation, as it is reducing $CO_2$ instead of oxidizing CO (Figure 2) – an energetically less favorable reaction. Mock et al. (2015) also observed that of the two highly transcribed CODH genes (CAETHG_1620-21 & 3005), the split gene formed a complex with acetyl-CoA synthase (ACS). Thus, the complex also catalyzes the most energetically unfavorable WLP reaction – the production of acetyl-CoA (Norman et al., 2018). Knocking-out this complex was lethal to *C. autoethanogenum* DSM 10061 growing autotrophically (Liew et al., 2016a). Additionally, a CAETHG_3005 knockout (KO) strain showed striking growth rate improvements when fermenting $CO_2$+$H_2$ compared to the wild type. However, this KO was detrimental to CO fermentation, where they observed a significant decrease in culture performance. Subsequently, Liew et al. (2016a) hypothesized that the extra CODH may act as a competitor for $CO_2$. In CO-containing fermentations by Valgepea et al. (2018) (high- and low-BC), it appeared that an enzyme after CODH in the WLP was limiting flux (Figures 2 & S3). Here, however, the WLP flux doubled with CO supplementation. Plausibly, an increase in CODH/ACS complex activity caused this. The mechanism for such an activity increase is unknown. We hypothesize there are four plausible options: increased enzyme turnover (better thermodynamics); increased enzyme concentration (upregulation); a combination of the first two; or isozyme-substrate competition – as hypothesized by Liew et al. (2016a). Confirming a mechanism, could make it possible to engineer the complex and gain further $CO_2$ or CO fermentation improvements. One means of testing this mechanism is to compare CO-supplemented $CO_2$+$H_2$ fermentation by wild type and CAETHG_3005 KO *C. autoethanogenum* on a multi-omics level.

#### Hydrogenase

Interestingly, we observed higher specific uptake of $H_2$ in $CO/CO_2/H_2$ compared to $CO+H_2$. Comparatively, HytABCDE (leq000001) production of reduced ferredoxin appears to drive AOR and Rnf complex (leq000004 and M002 respectively) in $CO/CO_2/H_2$[1]. Moreover, Nfn complex (leq000002) utilizes NADPH from HytABCDE, also to produce reduced ferredoxin. Higher ATP production, uses reduced ferredoxin *via* Rnf complex, and assists excretion of acetate from the cell, reducing product inhibition of the WLP and acidification of the cell (Figure 2) (Valgepea et al., 2018). Considering CO is a known inhibitor of hydrogenases (Adams, 1990; Bennett et al., 2000; De Lacey et al., 2007; Lubitz et al., 2007; Peters et al., 2015) a lower CO concentration may facilitate the increase in $H_2$ utilization compared to $CO+H_2$ fermentation. This is especially true for redox metabolism as the hydrogenase associated with carbon metabolism, HytABCDE/Fdh (rxn08518_c0) here (Wang et al., 2013), was found to have enhanced CO-tolerating properties in *Acetobacterium woodii* (Ceccaldi et al., 2017). These hypotheses agree with further relative and differential analysis of fluxes (Figure 2, and Tables S8 & S9). When normalized by carbon uptake, the differential change in HytABCDE flux between CO-containing fermentations and $CO/CO_2+H_2$[1] is greater than that of HytABCDE/Fdh. Yet, there is also ongoing discussion about CO activation of hydrogenases (Evans et al., 2016; Lamle et al., 2005), and CO involvement in hydrogenase maturation (CO ligand synthesis) (Bürstel et al., 2016; Fontecilla-Camps et al., 2007; Lubitz et al., 2014; Pagnier et al., 2016). Thus, there is still much to understand about how these enzymes function (Wittkamp et al., 2018), and their diversity (Peters et al., 2015).

#### AOR

For acetogenic, and especially $CO_2+H_2$ fermentation, redox metabolism is vital. The importance of AOR (leq000004) for *C. autoethanogenum*'s autotrophic redox metabolism was identified by Marcellin et al. (2016), and further characterized -enzymatically by Mock et al. (2015), -isozymatically by Liew et al. (2017), and -metabolically by Valgepea et al. (2018). When fermenting $CO_2+H_2$, it is believed that AOR activity is essential for ethanol production (Richter et al., 2016b), and dictates phenotypic differences between acetogens, for example *A. woodii* and *C. autoethanogenum* (Table 2)(Bertsch and Müller, 2015; Mock et al., 2015). As the AOR provides a reduction in acidification that also results in a industrially more valuable product, optimisation of this pathway could lead to industrially relevant shifts in carbon distributions.

### Supplementary Tables

Find Supplementary Tables – Large Tables in another supplementary file for larger tables.

### Table S1. Results summary

| Gas | Gas Composition | Gas flowrate | Stirrer Speed | Biological replicates | Dilution rate | Biomass | | Acetate | | Ethanol | |
|---|---|---|---|---|---|---|---|---|---|---|---|
| | (Ar to 100%) | mL/min | rpm | | $day^{-1}$ | gDCW/L | ± | g/L | ± | g/L | ± |
| CO | 60% CO | 50 | 510 | 4 | 1 | 0.47 | 0.02 | 2.12 | 0.18 | 0.63 | 0.05 |
| Syngas | 50 % CO, 20% $CO_2$, 20% $H_2$ | 50 | 500 | 2 | 1 | 0.48 | 0.04 | 4.35 | 0.12 | 0.61 | 0.06 |
| CO+$H_2$ | 15% CO, 45% $H_2$ | 50 | 650 | 4 | 1 | 0.46 | 0.04 | 0.69 | 0.07 | 4.46 | 0.41 |
| CO/$CO_2$/$H_2$ | 2% CO, 23% $CO_2$, 65% $H_2$ | 30 | 1200 | 2 | 1 | 0.34 | 0.02 | 5.03 | 0.34 | 4.79 | 0.43 |
| $CO_2$+$H_2$ | 23% $CO_2$, 67% $H_2$ | 32 | 500 | 3 | 0.5 | 0.18 | 0.02 | 2.51 | 0.42 | 2.36 | 0.25 |
| CO/$CO_2$/$H_2$ | 2% CO, 23% $CO_2$, 65% $H_2$ | 30 | 800 | 2 | 0.5 | 0.54 | 0.01 | 5.97 | 0.98 | 9.69 | 0.39 |

### Table S2. Flux ratios (mol%)

| Gas | Gas feed ratio | | | | Gas flux ratio | | | | Difference (Gas feed – Gas flux) | | | | Ethanol/substrate ratio (+acetate) | | | | |
|---|---|---|---|---|---|---|---|---|---|---|---|---|---|---|---|---|---|
| | $CO_2$/$H_2$ | CO/$H_2$ | CO+$CO_2$/$H_2$ | CO/$CO_2$ | $CO_2$/$H_2$ | CO/$H_2$ | CO+$CO_2$/$H_2$ | CO/$CO_2$ | $CO_2$/$H_2$ | CO/$H_2$ | CO+$CO_2$/$H_2$ | CO/$CO_2$ | EtOH/Ace | EtOH/$H_2$ | EtOH/CO | EtOH/$CO_2$ | EtOH/CO+$CO_2$ |
| CO | - | - | - | - | - | - | - | -1.33 | - | - | - | - | 0.41 | - | 0.06 | - | 0.06 |
| Syngas | 1.00 | 2.50 | 3.50 | 2.50 | -0.45 | 1.49 | 1.05 | -3.35 | 1.45 | 1.01 | 2.45 | 5.85 | 0.20 | 0.10 | 0.07 | - | 0.07 |
| CO+$H_2$ | - | 0.33 | 0.33 | - | -0.06 | 0.61 | 0.54 | -9.49 | - | -0.27 | -0.21 | - | 8.51 | 0.27 | 0.45 | - | 0.45 |
| CO/$CO_2$/$H_2$[1] | 0.35 | 0.03 | 0.38 | 0.09 | 0.36 | 0.03 | 0.40 | 0.09 | -0.01 | 0.00 | -0.01 | -0.01 | 1.26 | 0.12 | 3.43 | 0.32 | 0.30 |
| $CO_2$+$H_2$ | 0.34 | - | 0.34 | - | 0.41 | - | 0.41 | - | -0.06 | - | -0.06 | - | 1.24 | 0.12 | - | 0.31 | 0.31 |
| CO/$CO_2$/$H_2$[.5] | 0.35 | 0.03 | 0.38 | 0.09 | 0.34 | 0.02 | 0.37 | 0.07 | 0.01 | 0.01 | 0.02 | 0.02 | 2.15 | 0.13 | 5.46 | 0.37 | 0.34 |

The horizontal line through the middle of the table indicates where the data is from; above the line is data from (Valgepea et al., 2018), and below the line is novel data.

**Table S3. Flux values for Figure 1 (mmol/gDCW/day)**

| Gas | Value | Acetate | Ethanol | $CO_2$ | CO | $H_2$ |
|---|---|---|---|---|---|---|
| CO | AV | 76 | 31 | -394 | 523 | -14 |
| | SD | 5 | 1 | 6 | 12 | 1 |
| | SD% | 6% | 4% | 1% | 2% | -5% |
| Syngas | AV | 152 | 30 | -135 | 451 | 302 |
| | SD | 8 | 0 | 2 | 17 | 13 |
| | SD% | 5% | 0% | 1% | 4% | 4% |
| $CO+H_2$ | AV | 26 | 217 | -51 | 481 | 793 |
| | SD | 3 | 5 | 4 | 30 | 72 |
| | SD% | 12% | 2% | 9% | 6% | 9% |
| $CO/CO_2/H_2$[1] | AV | 243 | 306 | 944 | 89 | 2611 |
| | SD | 16 | 27 | 23 | 2 | 57 |
| | SD% | 7% | 9% | 2% | 2% | 2% |
| $CO_2+H_2$ | AV | 113 | 140 | 459 | 0 | 1130 |
| | SD | 9 | 12 | 77 | 0 | 158 |
| | SD% | 8% | 8% | 17% | 0% | 14% |
| $CO/CO_2/H_2$[.5] | AV | 91 | 197 | 538 | 36 | 1562 |
| | SD | 15 | 8 | 20 | 4 | 24 |
| | SD% | 16% | 4% | 4% | 11% | 2% |

**Table S4. Product carbon balance values for Figure 1 (mol%)**

| Gas | Value | Biomass | $CO_2$ | Acetate | Ethanol |
|---|---|---|---|---|---|
| CO | AV | 6.4 | 60.7 | 23.5 | 9.5 |
| | SD | 0.1 | 1.0 | 1.2 | 0.2 |
| | SD% | 1% | 2% | 5% | 3% |
| Syngas | AV | 7.7 | 25.0 | 56.3 | 11.1 |
| | SD | 0.3 | 0.5 | 1.1 | 0.3 |
| | SD% | 4% | 2% | 2% | 3% |
| $CO+H_2$ | AV | 7.2 | 8.9 | 8.4 | 75.5 |
| | SD | 0.1 | 0.8 | 0.2 | 0.9 |
| | SD% | 1% | 9% | 3% | 1% |
| $CO/CO_2/H_2$[1] | AV | 3.6 | - | 42.5 | 53.8 |
| | SD | 0.1 | - | 0.3 | 0.4 |
| | SD% | 2% | - | 1% | 1% |
| $CO_2+H_2$ | AV | 3.9 | - | 41.7 | 54.5 |
| | SD | 0.2 | - | 2.6 | 2.6 |
| | SD% | 6% | - | 6% | 5% |
| $CO/CO_2/H_2$[.5] | AV | 3.5 | - | 30.2 | 66.3 |
| | SD | 0.2 | - | 2.7 | 2.4 |
| | SD% | 7% | - | 9% | 4% |

AV – average, SD – standard deviation, SD% - standard deviation as a percentage of average flux.

**Table S5. Results summary for high-biomass comparison**

| Gas | Gas Composition | Gas flowrate | Stirrer Speed | Biological replicates | Dilution rate | Biomass | | Acetate | | Ethanol | |
|---|---|---|---|---|---|---|---|---|---|---|---|
| | (Ar to 100%) | mL/min | rpm | | $day^{-1}$ | gDCW/L | ± | g/L | ± | g/L | ± |
| CO | 60% CO | 50 | 665 | 4 | 1 | 1.43 | 0.08 | 6.28 | 0.43 | 3.88 | 0.15 |
| Syngas | 50 % CO, 20% $CO_2$, 20% $H_2$ | 50 | 650 | 2 | 1 | 1.36 | 0.06 | 7.86 | 0.14 | 5.43 | 0.33 |
| CO+H2 | 15% CO, 45% $H_2$ | 120 | 1000 | 4 | 1 | 1.45 | 0.04 | 3.84 | 0.33 | 11.55 | 0.41 |
| CO/CO2/H2 | 2% CO, 23% $CO_2$, 65% $H_2$ | 30 | 1200 | 2 | 1 | 0.34 | 0.02 | 5.03 | 0.34 | 4.79 | 0.43 |

The horizontal line through the middle of the table indicates where the data is from; above the line is data from (Valgepea et al., 2018), and below the line is novel data.

**Table S6. Volumetric flux and free energy balance of fermentations**

| Reaction | $\Delta G_R^0$, kJ/mol | Volumetric Productivity, mol/L/day | | | | | | | | |
|---|---|---|---|---|---|---|---|---|---|---|
| | | CO | | Syngas | | CO+$H_2$ | | $CO/CO_2/H_2$[1] | $CO_2$+$H_2$ | $CO/CO_2/H_2$[0.5] |
| | | LBC | HBC | LBC | HBC | LBC | HBC | | | |
| Biomass | 479 | 0.020 | 0.060 | 0.020 | 0.057 | 0.019 | 0.060 | 0.014 | 0.004 | 0.011 |
| Ethanol | 1300 | 0.014 | 0.089 | 0.014 | 0.123 | 0.099 | 0.274 | 0.105 | 0.026 | 0.107 |
| Acetate | 873 | 0.036 | 0.107 | 0.071 | 0.133 | 0.012 | 0.066 | 0.084 | 0.021 | 0.050 |
| Hydrogen | -230 | 0 | 0 | 0.146 | 0.387 | 0.363 | 1.026 | 0.900 | 0.209 | 0.849 |
| CO | -257 | 0.245 | 1.060 | 0.219 | 0.997 | 0.220 | 0.713 | 0.031 | 0 | 0.020 |
| $CO_2$ | 0 | -0.185 | -0.727 | -0.065 | -0.410 | -0.023 | -0.152 | 0.326 | 0.084 | 0.293 |
| **$\Delta \dot{G}_{OR}$, kJ/mol/day** | | **-3.6** | **-31.1** | **0.6** | **-42.2** | **8.5** | **23.6** | **2.0** | **5.4** | **-12.3** |

Fermentations from Valgepea et al. (2018) and this report. Biomass Gibb's free energy of formation was obtained from Liu et al. (2016). All other free energy values were obtained from (Chemistry Reference), see Free energy balance for details. LBC – low biomass concentration, HBC – high biomass concentration.

### Table S7. Metabolic highlights between $CO_2$-fermenting cultures

| Reaction | Reaction Flux mmol/gDCW/day | | | $CO_2$-Normalised Differential Flux mol/mol $CO_2$ | | |
|---|---|---|---|---|---|---|
| | $CO/CO_2/H_2^{1}$ | $CO_2$+$H_2$ | $CO/CO_2/H_2^{0.5}$ | $CO/CO_2/H_2^{1}$ - $CO_2$+$H_2$ | $CO/CO_2/H_2^{0.5}$ - $CO_2$+$H_2$ | 1-0.5 |
| **Exchange** | | | | | | |
| $CO_2$ | 42.64 | 21.00 | 22.43 | | | |
| CO | 3.71 | 0.00 | 1.51 | ↗ 0.09 | ↗ 0.07 | → 0.02 |
| $H_2$ | 114.81 | 54.21 | 63.13 | ↑ 0.11 | ↑ 0.23 | ↓ -0.12 |
| Acetate | 10.13 | 4.69 | 3.80 | → 0.01 | ↘ -0.05 | ↗ 0.07 |
| Ethanol | 12.56 | 5.83 | 8.06 | → 0.02 | ↗ 0.08 | ↘ -0.06 |
| **Central Metabolism** | | | | | | |
| CODH/ACS (Fdred) | -19.35 | -10.52 | -10.46 | → 0.05 | → 0.03 | → 0.01 |
| Tetrahydrofolate ligase (ATP) | -23.09 | -10.54 | -11.98 | → -0.04 | → -0.03 | → 0.00 |
| Methylene-THF oxidoreductase (NADPH) | -23.08 | -10.53 | -11.97 | → -0.04 | → -0.03 | → 0.00 |
| Methylene-THF reductase (Fdred) | 23.07 | 10.53 | 11.97 | → 0.04 | → 0.03 | → 0.00 |
| Acetyl-P Transferase (ATP) | 22.64 | 10.50 | 11.84 | → 0.03 | → 0.03 | → 0.00 |
| AOR (Fdred) | -12.54 | -5.82 | -8.04 | → -0.02 | ↘ -0.08 | ↗ 0.06 |
| Ethanol:NAD+ oxidoreductase (NADH) | -12.56 | -5.83 | -8.06 | → -0.02 | ↘ -0.08 | ↗ 0.06 |
| HytA-E (Fdred) | 45.86 | 21.84 | 25.57 | → 0.04 | ↑ 0.10 | ↘ -0.06 |
| Nfn Complex (Fdred) | 11.03 | 5.44 | 6.53 | → 0.00 | → 0.03 | → -0.03 |
| Rnf Complex (Fdred) | -47.79 | -21.51 | -25.51 | ↘ -0.10 | ↓ -0.11 | → 0.02 |
| ATPase (ATP) | 18.31 | 8.22 | 9.87 | → 0.04 | → 0.05 | → -0.01 |
| **Other** | | | | | | |
| PFOR (pyruvate) | 0.27 | 0.04 | 0.05 | → 0.00 | → 0.00 | → 0.00 |
| Total specific ATP production rate (ATP) | 40.96 | 18.72 | 21.71 | ↗ 0.07 | ↗ 0.08 | → 0.00 |
| H2O (H2O excretion) | 56.04 | 27.12 | 30.97 | → 0.02 | ↗ 0.09 | ↘ -0.07 |
| NGAM (ATP consumption) | 15.86 | 7.17 | 8.63 | → 0.03 | → 0.04 | → 0.00 |

Average specific fluxes were used for all comparisons. "Exchange" and "Central Metabolism" reactions match to Figure S4. See Flux data analysis for calculation details. Metabolites in brackets show forward direction of reaction (i.e. Rnf complex consumes reduced ferredoxin). Arrow scales are: green-up >= 0.1, yellow-diagonal-up >= 0.05, yellow-sideways >= -0.05, yellow-diagonal-down >= -0.1, red-down < -0.1. See Figure S1 for full metabolite and enzyme names and Figure S2 for full enzyme names and reaction IDs.

### Table S8. Highlights of comparative flux analysis

| Reaction | Flux, mmol/gDCW/h | | | | | | | | | Differential flux change compared to $CO/CO_2/H_2^1$, mmol/gDCW/h | | | | | | | |
|---|---|---|---|---|---|---|---|---|---|---|---|---|---|---|---|---|---|
| | a | b | c | d | e | f | g | h | i | a | b | c | d | e | f | h | i |
| ATP:acetate phosphotransferase | 4.42 | 5.64 | 7.52 | 7.80 | 9.95 | 9.75 | 22.64 | 10.50 | 11.84 | -18.22 | -17.00 | -15.12 | -14.84 | -12.69 | -12.90 | -12.14 | -10.80 |
| Proton transport | 5.80 | 7.91 | 13.49 | 12.35 | 10.54 | 11.51 | 32.19 | 14.54 | 15.09 | -26.39 | -24.28 | -18.70 | -19.83 | -21.65 | -20.68 | -17.65 | -17.09 |
| Methylene-THF reductase | 4.88 | 6.39 | 7.96 | 8.54 | 10.39 | 10.35 | 23.07 | 10.53 | 11.97 | -18.19 | -16.67 | -15.11 | -14.53 | -12.67 | -12.72 | -12.54 | -11.10 |
| Formate:THF ligase | 4.91 | 6.42 | 7.99 | 8.57 | 10.42 | 10.37 | 23.09 | 10.54 | 11.98 | -18.19 | -16.67 | -15.11 | -14.53 | -12.67 | -12.72 | -12.55 | -11.11 |
| 5,10-Methenyl-THF 5-hydrolase | -4.89 | -6.40 | -7.97 | -8.55 | -10.40 | -10.36 | -23.08 | -10.53 | -11.97 | 18.19 | 16.67 | 15.11 | 14.53 | 12.67 | 12.72 | 12.54 | 11.11 |
| ATP synthase | 7.45 | 9.50 | 8.33 | 10.39 | 11.92 | 11.91 | 18.31 | 8.22 | 9.87 | -10.87 | -8.82 | -9.98 | -7.92 | -6.40 | -6.40 | -10.10 | -8.45 |
| 5,10-methylene-THF oxidoreductase | -4.89 | -6.40 | -7.97 | -8.55 | -10.40 | -10.36 | -23.08 | -10.53 | -11.97 | 18.19 | 16.67 | 15.11 | 14.53 | 12.67 | 12.72 | 12.54 | 11.11 |
| Ethanol:NAD+ oxidoreductase | 1.29 | 2.58 | 1.24 | 3.78 | 8.99 | 7.90 | 12.56 | 5.83 | 8.06 | -11.27 | -9.98 | -11.32 | -8.78 | -3.57 | -4.66 | -6.73 | -4.51 |
| Acetyl-CoA acetyltransferase | 4.42 | 5.64 | 7.52 | 7.80 | 9.95 | 9.75 | 22.64 | 10.50 | 11.84 | -18.22 | -17.00 | -15.12 | -14.84 | -12.69 | -12.90 | -12.14 | -10.80 |
| NGAM | 3.03 | 5.55 | 5.96 | 7.72 | 9.56 | 9.34 | 15.86 | 7.17 | 8.63 | -12.83 | -10.31 | -9.90 | -8.13 | -6.29 | -6.51 | -8.69 | -7.23 |
| H2O transport | 11.65 | 15.79 | 3.92 | 8.44 | -7.53 | -4.27 | -56.04 | -27.12 | -30.97 | 67.69 | 71.83 | 59.96 | 64.48 | 48.51 | 51.78 | 28.92 | 25.07 |
| CO2 transport | -16.42 | -21.12 | -5.61 | -12.57 | -2.11 | -4.39 | 42.64 | 21.00 | 22.43 | -59.07 | -63.76 | -48.25 | -55.21 | -44.75 | -47.03 | -21.65 | -20.22 |
| CO transport | -26.44 | -34.32 | -21.91 | -30.57 | -23.19 | -25.54 | -3.72 | 0.00 | -1.51 | -22.72 | -30.60 | -18.19 | -26.85 | -19.48 | -21.83 | 3.72 | 2.21 |
| CO dehydrogenase | 21.57 | 27.93 | 13.95 | 22.03 | 12.80 | 15.20 | -19.35 | -10.52 | -10.46 | 40.91 | 47.28 | 33.30 | 41.38 | 32.15 | 34.55 | 8.82 | 8.89 |
| CODH/ACS | 4.87 | 6.39 | 7.95 | 8.53 | 10.39 | 10.34 | 23.06 | 10.52 | 11.96 | -18.19 | -16.67 | -15.11 | -14.53 | -12.67 | -12.72 | -12.54 | -11.10 |
| 5-Methyl-THF methyltransferase | 4.87 | 6.39 | 7.95 | 8.53 | 10.39 | 10.34 | 23.06 | 10.52 | 11.96 | -18.19 | -16.67 | -15.11 | -14.53 | -12.67 | -12.72 | -12.54 | -11.10 |
| H2 transport | 0.58 | 0.49 | -13.47 | -12.99 | -37.39 | -32.84 | -114.81 | -54.21 | -63.13 | 115.38 | 115.30 | 101.34 | 101.82 | 77.41 | 81.97 | 60.59 | 51.68 |
| Ethanol transport | 1.29 | 2.58 | 1.24 | 3.78 | 8.99 | 7.90 | 12.56 | 5.83 | 8.06 | -11.27 | -9.98 | -11.32 | -8.78 | -3.57 | -4.66 | -6.73 | -4.51 |
| Rnf complex | 16.64 | 21.63 | 20.28 | 24.70 | 28.68 | 28.63 | 47.79 | 21.51 | 25.51 | -31.15 | -26.16 | -27.51 | -23.09 | -19.11 | -19.16 | -26.28 | -22.27 |
| Acetate transport | 3.18 | 3.11 | 6.32 | 4.06 | 1.00 | 1.89 | 10.13 | 4.69 | 3.80 | -6.95 | -7.02 | -3.81 | -6.07 | -9.12 | -8.24 | -5.43 | -6.32 |
| HytABCDE | -0.29 | -0.25 | 2.72 | 1.98 | 13.48 | 11.23 | 45.86 | 21.84 | 25.57 | -46.14 | -46.10 | -43.13 | -43.88 | -32.37 | -34.63 | -24.02 | -20.28 |
| Nfn complex | 5.48 | 6.08 | 2.84 | 3.50 | -1.27 | -0.18 | -11.03 | -5.44 | -6.53 | 16.51 | 17.11 | 13.86 | 14.52 | 9.76 | 10.84 | 5.58 | 4.50 |
| Formate hydrogen-lyase | 0.00 | 0.00 | 8.02 | 9.02 | 10.43 | 10.37 | 23.09 | 10.54 | 11.98 | -23.09 | -23.09 | -15.07 | -14.07 | -12.67 | -12.72 | -12.55 | -11.11 |
| AOR | 1.27 | 2.56 | 1.22 | 3.76 | 8.97 | 7.88 | 12.54 | 5.82 | 8.04 | -11.27 | -9.98 | -11.32 | -8.78 | -3.57 | -4.66 | -6.72 | -4.49 |
| PFOR | -0.31 | -0.60 | -0.29 | -0.59 | -0.30 | -0.45 | -0.27 | -0.04 | -0.05 | -0.04 | -0.33 | -0.02 | -0.32 | -0.02 | -0.17 | 0.32 | 0.22 |

### Table S9. Highlights of comparative flux analysis normalised by carbon uptake

| Reaction | Absolute-relative flux change normalised by carbon uptake and compared to $CO/CO_2/H_2^1$, mol/mol | | | | | | | | Differential flux change normalised by carbon uptake and compared to $CO/CO_2/H_2^1$, mol/mol | | | | | | | |
|---|---|---|---|---|---|---|---|---|---|---|---|---|---|---|---|---|
| | a | b | c | d | e | f | h | i | a | b | c | d | e | f | h | i |
| ATP:acetate phosphotransferase | 0.34 | 0.34 | 0.70 | 0.52 | 0.88 | 0.78 | 1.02 | 1.01 | 0.32 | 0.32 | 0.15 | 0.23 | 0.06 | 0.11 | -0.01 | -0.01 |
| Proton transport | 0.32 | 0.33 | 0.89 | 0.58 | 0.65 | 0.65 | 1.00 | 0.91 | 0.47 | 0.46 | 0.08 | 0.29 | 0.24 | 0.24 | 0.00 | 0.06 |
| Methylene-THF reductase | 0.37 | 0.37 | 0.73 | 0.56 | 0.90 | 0.81 | 1.01 | 1.00 | 0.31 | 0.31 | 0.13 | 0.22 | 0.05 | 0.09 | 0.00 | 0.00 |
| Formate:THF ligase | 0.37 | 0.38 | 0.73 | 0.56 | 0.90 | 0.82 | 1.01 | 1.00 | 0.31 | 0.31 | 0.13 | 0.22 | 0.05 | 0.09 | 0.00 | 0.00 |
| 5,10-Methenyl-THF 5-hydrolase | 0.37 | 0.37 | 0.73 | 0.56 | 0.90 | 0.81 | 1.01 | 1.00 | -0.31 | -0.31 | -0.13 | -0.22 | -0.05 | -0.09 | 0.00 | 0.00 |
| ATP synthase | 0.71 | 0.70 | 0.96 | 0.86 | 1.30 | 1.18 | 0.99 | 1.04 | 0.11 | 0.12 | 0.01 | 0.05 | -0.12 | -0.07 | 0.00 | -0.02 |
| 5,10-methylene-THF oxidoreductase | 0.37 | 0.37 | 0.73 | 0.56 | 0.90 | 0.81 | 1.01 | 1.00 | -0.31 | -0.31 | -0.13 | -0.22 | -0.05 | -0.09 | 0.00 | 0.00 |
| Ethanol:NAD+ oxidoreductase | 0.18 | 0.28 | 0.21 | 0.46 | 1.43 | 1.14 | 1.02 | 1.24 | 0.22 | 0.20 | 0.21 | 0.15 | -0.12 | -0.04 | -0.01 | -0.07 |
| Acetyl-CoA acetyltransferase | 0.34 | 0.34 | 0.70 | 0.52 | 0.88 | 0.78 | 1.02 | 1.01 | 0.32 | 0.32 | 0.15 | 0.23 | 0.06 | 0.11 | -0.01 | -0.01 |
| NGAM | 0.34 | 0.47 | 0.79 | 0.74 | 1.21 | 1.07 | 1.00 | 1.05 | 0.23 | 0.18 | 0.07 | 0.09 | -0.07 | -0.02 | 0.00 | -0.02 |
| H2O transport | 0.36 | 0.38 | 0.15 | 0.23 | 0.27 | 0.14 | 1.07 | 1.07 | -1.65 | -1.67 | -1.39 | -1.49 | -0.88 | -1.04 | 0.08 | 0.09 |
| CO2 transport | 0.68 | 0.67 | 0.28 | 0.45 | 0.10 | 0.19 | 1.09 | 1.02 | 1.54 | 1.54 | 1.18 | 1.33 | 1.01 | 1.09 | -0.08 | -0.02 |
| CO transport | 12.48 | 12.48 | 12.48 | 12.48 | 12.48 | 12.48 | 0.00 | 0.78 | 0.92 | 0.92 | 0.92 | 0.92 | 0.92 | 0.92 | -0.08 | -0.02 |
| CO dehydrogenase | 1.95 | 1.95 | 1.53 | 1.73 | 1.32 | 1.43 | 1.20 | 1.05 | -1.23 | -1.23 | -1.05 | -1.14 | -0.97 | -1.01 | 0.08 | 0.02 |
| CODH/ACS | 0.37 | 0.37 | 0.73 | 0.56 | 0.90 | 0.81 | 1.01 | 1.00 | 0.31 | 0.31 | 0.13 | 0.22 | 0.05 | 0.09 | 0.00 | 0.00 |
| 5-Methyl-THF methyltransferase | 0.37 | 0.37 | 0.73 | 0.56 | 0.90 | 0.81 | 1.01 | 1.00 | 0.31 | 0.31 | 0.13 | 0.22 | 0.05 | 0.09 | 0.00 | 0.00 |
| H2 transport | 0.01 | 0.01 | 0.25 | 0.17 | 0.65 | 0.52 | 1.04 | 1.07 | -2.50 | -2.49 | -1.86 | -2.05 | -0.86 | -1.19 | 0.11 | 0.16 |
| Ethanol transport | 0.18 | 0.28 | 0.21 | 0.46 | 1.43 | 1.14 | 1.02 | 1.24 | 0.22 | 0.20 | 0.21 | 0.15 | -0.12 | -0.04 | -0.01 | -0.07 |
| Rnf complex | 0.61 | 0.61 | 0.90 | 0.78 | 1.20 | 1.09 | 0.99 | 1.03 | 0.40 | 0.40 | 0.10 | 0.22 | -0.21 | -0.09 | 0.01 | -0.04 |
| Acetate transport | 0.55 | 0.41 | 1.32 | 0.61 | 0.20 | 0.34 | 1.02 | 0.73 | 0.10 | 0.13 | -0.07 | 0.09 | 0.18 | 0.14 | -0.01 | 0.06 |
| HytABCDE | 0.01 | 0.01 | 0.13 | 0.07 | 0.59 | 0.44 | 1.05 | 1.08 | 1.00 | 1.00 | 0.86 | 0.92 | 0.41 | 0.55 | -0.05 | -0.08 |
| Nfn complex | 0.87 | 0.75 | 0.54 | 0.48 | 0.23 | 0.03 | 1.09 | 1.15 | -0.45 | -0.42 | -0.37 | -0.35 | -0.18 | -0.23 | 0.02 | 0.03 |
| Formate hydrogen-lyase | 0.00 | 0.00 | 0.74 | 0.59 | 0.90 | 0.82 | 1.01 | 1.00 | 0.50 | 0.50 | 0.13 | 0.20 | 0.05 | 0.09 | 0.00 | 0.00 |
| AOR | 0.18 | 0.28 | 0.21 | 0.45 | 1.43 | 1.14 | 1.02 | 1.24 | 0.22 | 0.20 | 0.21 | 0.15 | -0.12 | -0.04 | -0.01 | -0.07 |
| PFOR | 1.99 | 2.97 | 2.26 | 3.27 | 2.15 | 2.96 | 0.35 | 0.37 | 0.01 | 0.01 | 0.01 | 0.01 | 0.01 | 0.01 | -0.01 | 0.00 |

Average specific fluxes were used for all comparisons. For full analysis see Figure S2. Figure S2 also contains full description and ID for noted reactions. a & b – CO low & high biomass, c & d syngas low & high biomass, e & f – CO+$H_2$ low & high biomass, g – $CO/CO_2/H_2^1$, h – $CO_2$+$H_2$, i – $CO/CO_2/H_2^{0.5}$. Division and subtraction by the corresponding flux of $CO/CO_2/H_2^1$, respectively, produced the relative and differential analysis. The absolute-relative change was used here (compared to non-absolute in Figure S2) to show change as an increase (green), decrease (red), and no change (black) in flux magnitude. Scales (red to green): (top left) -30 to 30, (top right) -50 to 50, (bottom left) 0 to 2, (bottom right) -1 to 1. Positive/negative values do not implicitly lead to higher/lower activity, respectively. These comparisons were used only as an indication of where change was occurring. To ascertain increase/decrease effect precisely see Supplementary Tables S10-12 for flux data.

### Supplementary Figures

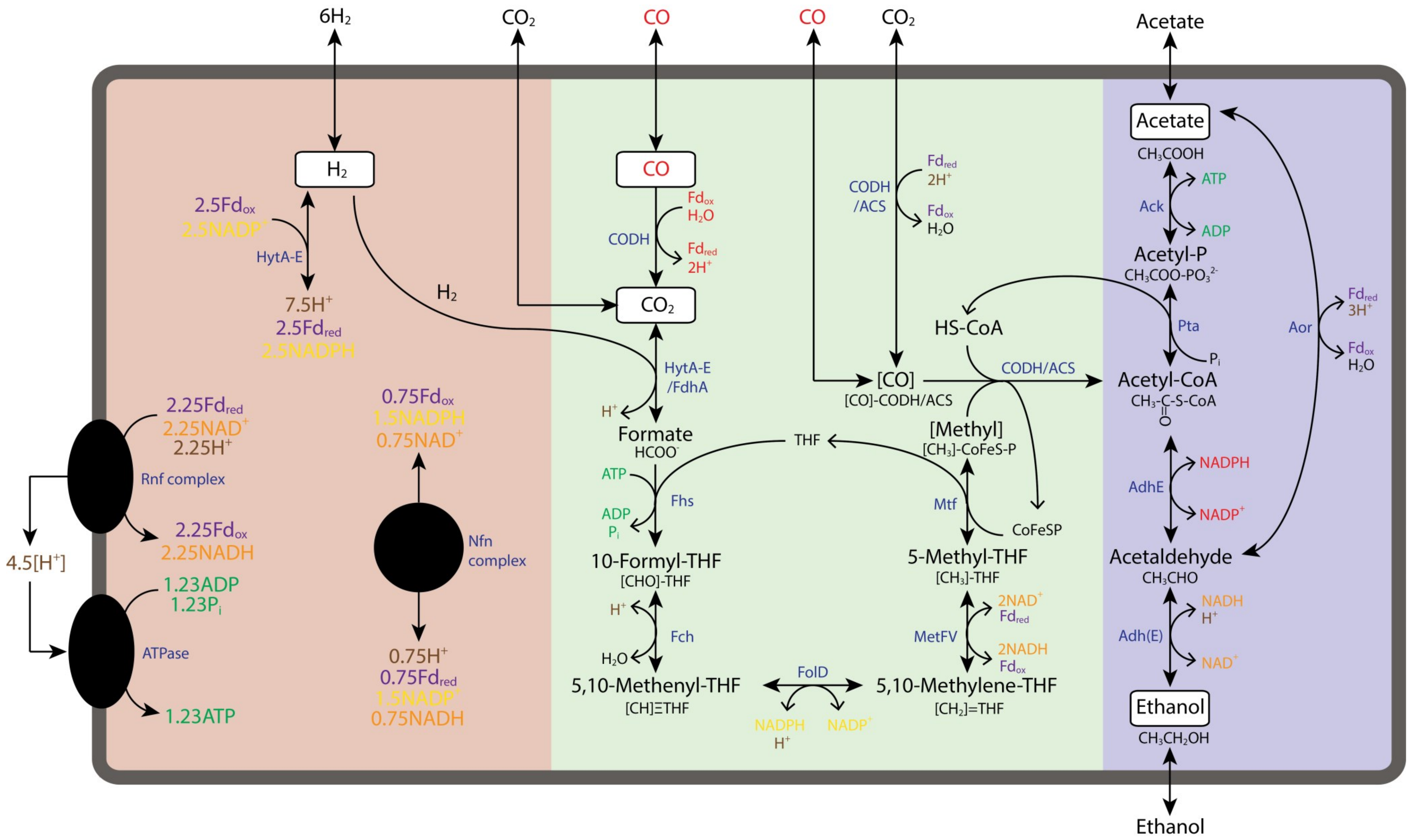
6H2
CO2
CO
CO
CO2
Acetate
H2
CO
Acetate
CH3COOH
2.5Fdox
2.5NADP+
HytA-E
7.5H+
2.5Fdred
2.5NADPH
H2
Fdox
H2O
CODH
Fdred
2H+
CO2
CODH
/ACS
Fdred
2H+
Fdox
H2O
Ack
ATP
ADP
Acetyl-P
CH3COO-PO3 2-
Pta
Pi
HS-CoA
Aor
Fdred
3H+
Fdox
H2O
HytA-E
/FdhA
H+
[CO]
[CO]-CODH/ACS
CODH/ACS
Acetyl-CoA
CH3-C-S-CoA
O
2.25Fdred
2.25NAD+
2.25H+
Rnf complex
2.25Fdox
2.25NADH
0.75Fdox
1.5NADPH
0.75NAD+
Nfn
complex
Formate
HCOO-
ATP
ADP
Pi
Fhs
THF
[Methyl]
[CH3]-CoFeS-P
Mtf
CoFeSP
AdhE
NADPH
NADP+
4.5[H+]
10-Formyl-THF
[CHO]-THF
5-Methyl-THF
[CH3]-THF
Acetaldehyde
CH3CHO
1.23ADP
1.23Pi
ATPase
1.23ATP
0.75H+
0.75Fdred
1.5NADP+
0.75NADH
H+
H2O
Fch
MetFV
2NAD+
Fdred
2NADH
Fdox
Adh(E)
NADH
H+
NAD+
5,10-Methenyl-THF
[CH]≡THF
FolD
NADPH
H+
NADP+
5,10-Methylene-THF
[CH2]=THF
Ethanol
CH3CH2OH
Ethanol

**Figure. S1.** Central metabolic pathways of *Clostridium autoethanogenum* when fermenting $CO_2$+$H_2$ to ethanol. Here, ferredoxin was assumed to be the electron acceptor of the reaction catalysed by MetFV and ethanol production was assumed to occur via the AOR. The overall reaction was $2CO_2 + 6H_2 + 1.23ADP + 1.23P_i \rightarrow CH_3CH_2OH + 3H_2O + 1.23ATP$. Reactions involving free CO and the reaction facilitated by AdhE (acetyl-CoA to acetaldehyde) are not active. Arrows show the reaction direction. **Background**: red – energetic/redox pathways, green – WLP, blue – product pathways. **Writing**: reaction not present (red), $Fd_{ox}$ and $Fd_{red}$ - oxidised and reduced ferredoxin (purple), NADH and $NAD^+$ – protonated and deprotonated nicotinamide adenine dinucleotide (orange), NADPH and $NADP^+$ - protonated and deprotonated NAD phosphate (yellow), ATP and ADP + $P_i$ – adenosine triphosphate and precursors (green), $H^+$ – protons (brown), THF – tetrahydrofolate, **enzymes** (blue): HytA-E – electron bifurcating, NADP - and ferredoxin-dependent [FeFe]-hydrogenase, HytA-E/FdhA – HytA-E in complex with selenium- and tungsten-dependent formate dehydrogenase, CODH – carbon monoxide dehydrogenase, CODH/ACS – CODH complex with acetyl CoA synthase, Rnf – $Fd_{red}$:$NAD^+$ oxidoreductase, Nfn – ferredoxin-dependent transhydrogenase, ATPase – ATP synthase, Fhs – formate:THF ligase, Fch – 5,10-methenyl-THF 5-hydrolase, FolD – methylene-THF dehydrogenase, MetFV – methylene-THF reductase, Mtf - methyl-THF:CoFeSP transferase, Pta – phosphotransacetylase, Ack – acetate kinase, AdhE - bifunctional alcohol dehydrogenase/acetaldehyde dehydrogenase, Adh - alcohol dehydrogenase, Aor - acetylaldehyde:ferredoxin oxidoreductase.

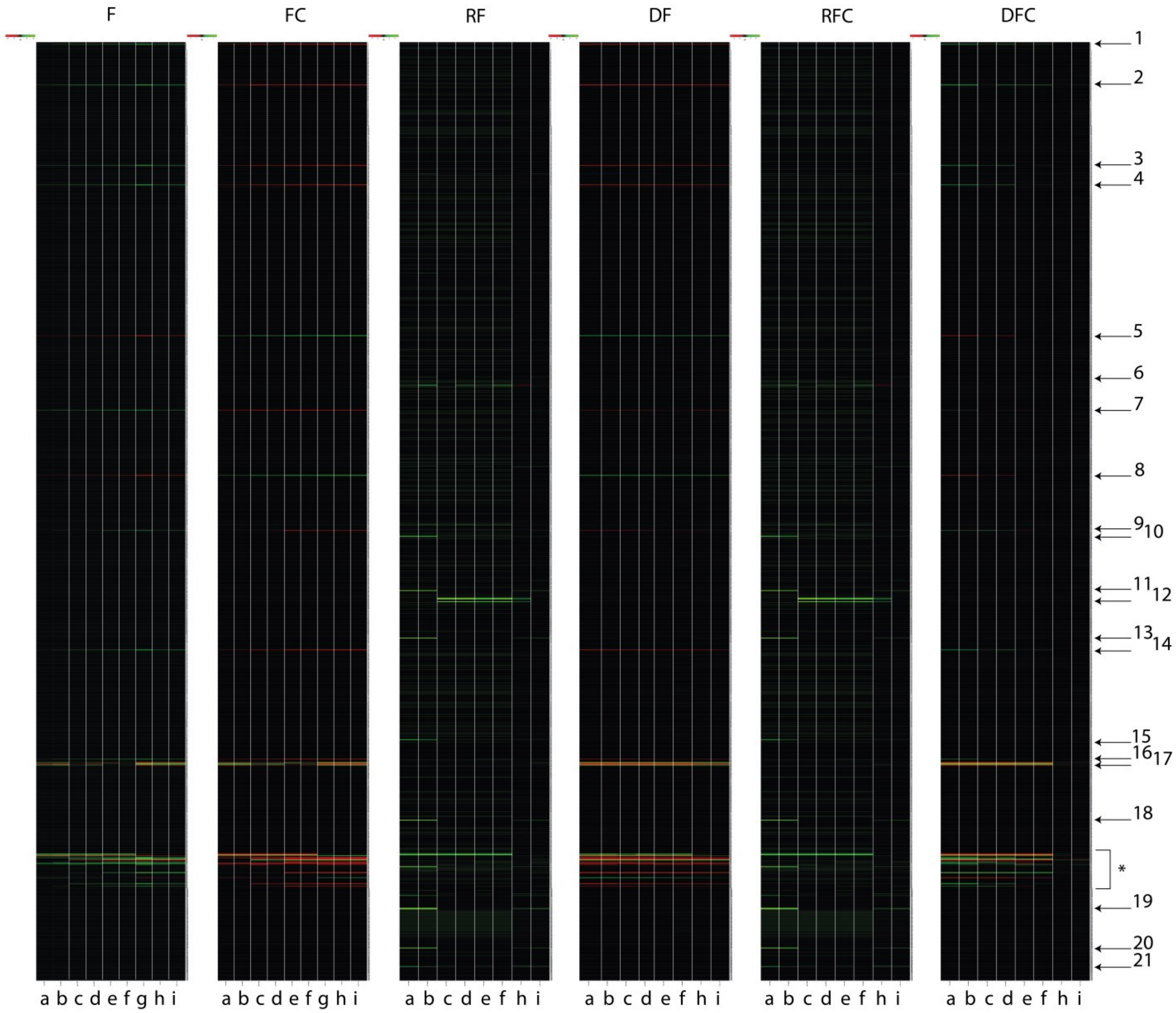


**Figure S2.** Initial comparative flux analysis of entire FBA datasets with heatmaps. All fluxes are an average of the respective steady-state biological replicates, with further analysis also completed using these averages. F – flux, FC – flux normalised by carbon uptake, RF – relative flux, DF – differential flux, RFC – relative flux normalised by carbon uptake, DFC – differential flux normalised by carbon uptake. a & b – CO low & high biomass, c & d – syngas low & high biomass, e & f – CO+$H_2$ low & high biomass, g – $CO/CO_2/H_2^1$, h – $CO_2$+$H_2$, i – $CO/CO_2/H_2^{0.5}$. Division and subtraction by the corresponding flux of $CO/CO_2/H_2^1$ produced the relative and differential analysis, respectively. Colour scales are roughly (red to green respectively): -100 to 100 for F & DF, -2 to 2 for FC & DFC, -25 to 25 RF, and -50 to 50 for RFC. Results from Valgepea et al. (2018) are also displayed, the conditions of these fermentations are summarized in Table S1 & S7. Carbon uptake was a negative value, hence the directionality switch (e.g. F to FC). Reactions are displayed in the same order as in the Supplementary Tables S10-12. See Tables S8 & S9 for highlights of the heatmap. Heatmaps were produced using Heatmapper (Babicki et al., 2016).

Reactions noted in the heatmap: 1 – acetate phosphotransferase (rxn00225_c0); 2 – proton transport (cpd00067_ext_b); 3 – NADH-dependent, electron-bifurcating, methylene-THF reductase (rxn00910_c0); 4 – formate:THF ligase (rxn00690_c0); 5 – 5,10-methenyl-THF 5-hydrolase (rxn01211_c0); 6 – ammonia diffusion (rxn05466_c0 ); 7 – ATP synthase (rxn10042_c0); 8 – 5,10-methylene-THF:NADP$^+$ oxidoreductase (rxn00907_c0); 9 – ethanol:NAD$^+$ oxidoreductase (rxn00543_c0); 10 – ATP:AMP phosphotransferase (rxn00097_c0); 11 – sulfate transport via proton symport (rxn05651_c0);12 – lactate transport (rxn05602_c0); 13 – thioredoxin reductase (rxn05289_c0); 14 – acytl-CoA:orthophosphate acetyltransferase (rxn00173_c0); 15 – pyrophosphate phosphohydrolase (rxn00001_c0); 16 – NGAM (ATP phosphohydrolase [protein-secreting]) (rxn00062_c0); 17 – water transport (rxn05319_c0) and $CO_2$ transport (rxn05467_c0); 18 – hydrogen-sulfide:NADP+ oxidoreductase (rxn00623_c0); * – CO transport (rxn10480_c0), CO dehydrogenase (rxn07189_c0), formate dehydrogenase (rxn00103_c0), CO dehydrogenase/acetyl-CoA synthase (rxn12080_c0), 5-methyl-THF:corrinoid Co-methyltransferase (rxn06149_c0), hydrogen transport (rxn10542_c0), ethanol transport (rxn09683_c0), Rnf complex (M002), acetate transport (rxn05488_c0), NADP-dependent electron-bifurcating [FeFe]-hydrogenase (HytABCDE) (leq000001), Nfn complex (leq000002), formate dehydrogenase/HytABCDE complex (rxn08518_c0), acetaldehyde:ferredoxin oxidoreductase (leq000004); 19 – sulfate adenyltransferase (rxn09240_c0); 20 – sulfate transport (cpd00048_ext_b); 21 – hydrogen sulphide transport (H2STrans_c0).

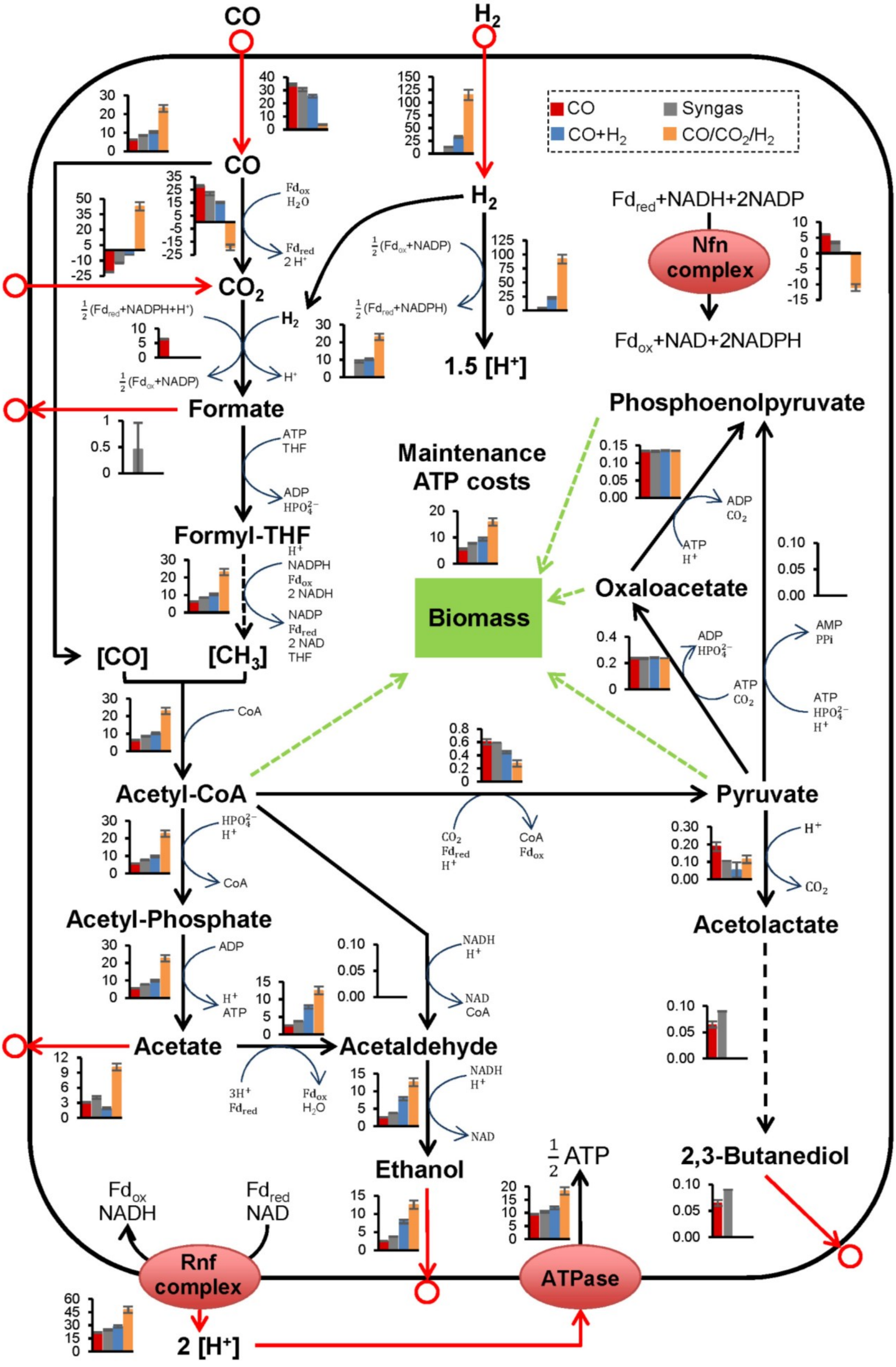
CO
H2
CO
Syngas
CO+H2
CO/CO2/H2
CO
H2
Fdred+NADH+2NADP
Nfn complex
Fdox+NAD+2NADPH
CO2
1.5 [H+]
Formate
Formyl-THF
Phosphoenolpyruvate
Maintenance ATP costs
Biomass
Oxaloacetate
[CO]
[CH3]
Acetyl-CoA
Pyruvate
Acetyl-Phosphate
Acetolactate
Acetate
Acetaldehyde
Ethanol
2,3-Butanediol
1/2 ATP
Fdox NADH
Fdred NAD
Rnf complex
ATPase
2 [H+]

**Figure S3.** Predictions of central metabolic pathway fluxes for high-biomass, autotrophic fermentations of *Clostridium autoethanogenum* using iCLAU786, flux balance analysis, and chemostat data. Results from Valgepea et al. (2018) are also displayed, the conditions of these fermentations are summarized in Table S5. Fluxes (mmol/gDCW/h) are represented as the average ± standard deviation between biological replicates. Number of biological replicates, and detailed gas composition for each fermentation are available in Table S5. Arrows show the direction of calculated fluxes; red arrows denote uptake or secretion, dashed arrows denote a series of reactions. Brackets denote metabolites bound by an enzyme. Refer to Figure S1 & S2 for enzyme involvement, metabolite abbreviations, and reaction IDs, and Supplementary Tables S10-12 for complete flux balance analysis datasets.

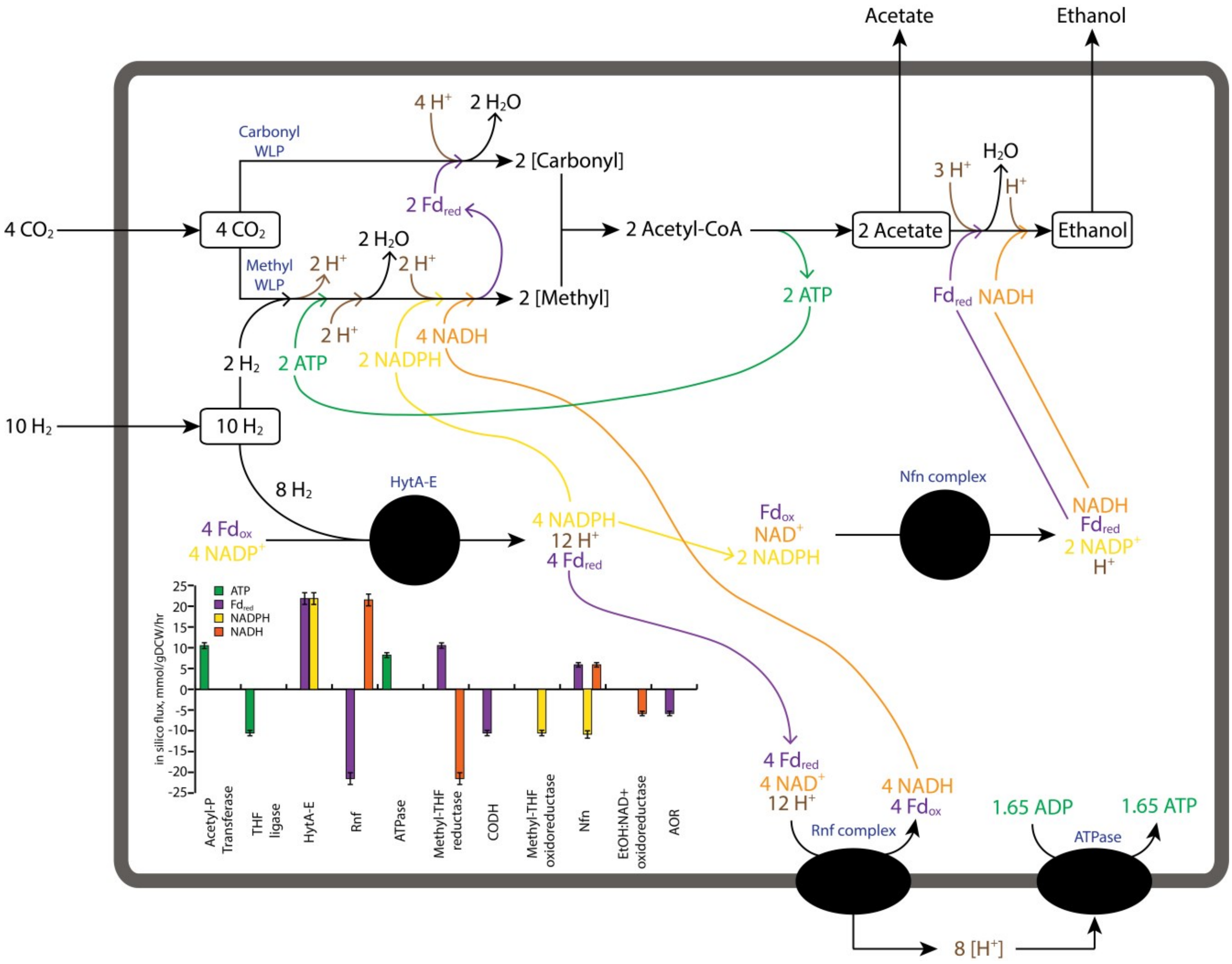


**Figure S4.** 1:1 mapping of WLP energetic cofactors indicated by flux balance analysis of *C. autoethanogenum* fermenting $CO_2$+$H_2$. The approximate experimental stoichiometry was $4CO_2 + 10H_2 + 1.65\ ADP + 1.65\ P_i \longrightarrow CH_3COOH + CH_3CH_2OH + 5H_2O + 1.65ATP$; where 1 "stoichiometric reaction mol" (figure pathways) = ~5 mmol/gDCW/hr (graph insert from FBA analysis). Refer to Table S7 for flux data shown here and comparison to CO-supplemented $CO_2$+$H_2$ fermentations, Figure S1 & S2 for enzyme involvement, metabolite abbreviations, and reaction IDs, and Supplementary Tables S10-12 for complete flux balance analysis datasets.
